# Fern cell walls and the evolution of arabinogalactan-proteins in streptophytes

**DOI:** 10.1101/2022.12.15.520549

**Authors:** Kim-Kristine Mueller, Lukas Pfeifer, Lina Schuldt, Péter Szövényi, Sophie de Vries, Jan de Vries, Kim L. Johnson, Birgit Classen

## Abstract

Significant changes have occurred in plant cell wall composition during evolution and diversification of tracheophytes. As the sister lineage to seed plants, knowledge on the cell wall of ferns is key to track evolutionary changes across tracheophytes and to understand seed plant-specific evolutionary innovations. Fern cell wall composition is not fully understood, including limited knowledge of glycoproteins such as the fern arabinogalactan-proteins (AGPs). Here, we characterize the AGPs from the leptosporangiate fern genera *Azolla*, *Salvinia* and *Ceratopteris*. The carbohydrate moiety of seed plant AGPs consists of a galactan backbone including mainly 1,3- and 1,3,6-linked pyranosidic galactose, which is conserved across the investigated fern AGPs. Yet, unlike AGPs of angiosperms, those of ferns contained the unusual sugar 3-*O*-methylrhamnose. Besides terminal furanosidic Ara (Ara*f*), the main linkage type of Ara*f* in the ferns was 1,2-linked Ara*f*, whereas in seed plants 1,5-linked Ara*f* is often dominating. Antibodies directed against carbohydrate epitopes of AGPs supported the structural differences between AGPs of ferns and seed plants. Comparison of AGP linkage types across the streptophyte lineage showed that angiosperms have rather conserved monosaccharide linkage types; by contrast bryophytes, ferns and gymnosperms showed more variability. Phylogenetic analyses of glycosyltransferases involved in AGP biosynthesis and bioinformatic search for AGP protein backbones revealed a versatile genetic toolkit for AGP complexity in ferns. Our data reveal important differences across AGP diversity which functional significance is unknown. This diversity sheds light on the evolution of the hallmark feature of tracheophytes: their elaborate cell walls.

**SIGNIFICANCE STATEMENT:** Ferns are the sister lineage of seed plants and key to understanding plant evolution. To understand ferns’ unique cell walls, we analysed arabinogalactan-proteins from the fern genera *Azolla*, *Salvinia* and *Ceratopteris*. Comparison of AGP structures throughout the streptophyte lineage reveals special features in relation to systematic positions and proposes a trend to more hydrophilic AGPs in course of evolution. Through comparative genomic analyses, we pinpoint the potential genetic players for this diversity in cell walls.

## INTRODUCTION

One of the most important steps in the evolution of life on Earth happened around 500 million years ago (mya), when descendents of streptophyte algae colonized the terrestrial habitat (e.g. Kenrick and Crane, 1997; Becker and Marin, 2009; Delwiche and Cooper, 2015; Bowman *et al*., 2017; Harrison, 2017; de Vries and Archibald, 2018). By the end of the Devonian (360 mya), the extant lineages of land plants comprising bryophytes, lycophytes, monilophytes (=ferns), and spermatophytes have evolved and the major organ and tissue types known from extant plants (e.g. vasculature, roots, leaves, seeds, wood, secondary growth) were already present (Bowman, 2013). Vascular plants (tracheophytes) comprise lycophytes, ferns and seed plants, and are characterized by features like tracheids and sieve elements for water and nutrient transport and a highly structured and dominant sporophyte. Lycophytes separated from all other living vascular plant lineages already in the nearly-mid Devonian (approx. 400 mya; Banks *et al*., 2011; Pryer *et al*., 2004), ferns first appeared in the early Carboniferous (Galtier and Scott, 1985). They comprise around 12,000 extant species belonging to the 5 lineages Equisetales, Psilotales, Ophioglossales, Maratialles and the dominant group of leptosporangiate ferns (Pryer *et al*., 2004; PPG 2016; Nitta *et al*., 2022). Ferns form the sister lineage to seed plants; to understand the evolutionary history of innovations specific to seed-plants, knowledge of this group is necessary to infer (i) the characteristics of the last common ancestor (LCA) shared by ferns and seed-plants and (ii) define the evolutionary trajectory of seed-plant specific characteristics (Rensing, 2017).

Ferns are known for possessing large genomes with numerous chromosomes. To date, genome sequence of the hetereosporous ferns *Azolla filiculoides* and *Salvinia cucullata*, the homosporous ferns *Ceratopteris richardii*, *Adiantum capillus-veneris*, as well as the tree fern *Alsophila spinulosa* are publicly available (Li *et al*., 2018; Marchant *et al*., 2022; Fang *et al*., 2022; Huang *et al*., 2022). *Azolla* and *Salvinia* are floating aquatic ferns with high growth rates and the potential to be significant carbon sinks. Data from the Arctic Ocean revealed that around 50 mya, an abundance of *Azolla* characterized an 800,000-year interval called the „Azolla event“ which possibly had a role in global cooling by sequestering atmospheric carbon dioxide (Brinkhuis *et al*., 2006; Speelman *et al*., 2009). *Azolla* is further remarkable due to its obligate symbiosis with the N-fixing cyanobacterium *Nostoc azollae* in specialized leaf cavities (Li *et al*., 2018; de Vries et al., 2018; de Vries and de Vries, 2022). *Ceratopteris richaridii* is also adapted to water and is known as the genetically tractable fern model organism („C-Fern“ Plackett *et al*., 2015) and as a teaching tool in biology (Renzaglia and Warne, 1995). *Azolla*, *Salvinia* and *Ceratopteris* all belong to the leptosporangiate ferns, which account for approximately 80 % of non-flowering vascular plant species (Plackett *et al*., 2015). Among them *Ceratopteris* was suggested as the model for research on adaptive cell wall modification in vascular plants (Leroux *et al*., 2013a).

Plant cell walls are important at various levels of plant morphology and as such are expected to have changed during evolution (Sørensen *et al*., 2011). According to current knowledge, fern cell walls share basic features with those of seed plants, e.g. the ocurrence of cellulose, hemicelluloses and pectic polysaccharides (Leroux *et al*., 2013a; Matsunaga *et al*., 2004; Popper and Fry, 2004; Popper, 2008). However, differences exist, e.g. in lignin structures: Whereas secondary cell walls of lycophytes and angiosperms are reinforced with lignin containing mainly syringyl monomers, lignin of ferns and gymnosperms is derived primarily from guaiacyl monomers (Weng *et al*., 2008). Besides polysaccharides and lignin, hydroxyproline-rich-glycoproteins (HRGPs) are also important components of plant cell walls. They are broadly classified into the three groups arabinogalactan-proteins (AGPs), extensins and proline-rich proteins (PRPs) however in reality a continuum exists (Johnson *et al*., 2018). PRPs are only minimally glycosylated, extensins possess a glycan part of around 50 % and AGPs consist of up to 90 % polysaccharides. In AGPs, several arabinogalactan (AG) moieties are covalently linked to the protein *via* hydroxyproline. The characteristic structure of the AG in angiosperms is a backbone of 1,3-linked β-d-Gal*p*, branched at position 6 to 1,6-linked β-d-Gal*p* side chains, which are substituted with α-l-Ara*f* and often also terminal β-d-Glc*p*A (for review see Ma *et al*., 2018; Strasser *et al*., 2021). AGPs are key constituents of seed plants extracellular matrix and are involved in several processes such as cell growth, cell proliferation, pattern formation, sexual reproduction and plant-microbe interactions (for review see Seifert and Roberts, 2007; Ma *et al*., 2018).

A unique feature of AGPs is their ability to precipitate with red-coloured Yariv phenylglycosides, e.g. the β-glucosyl Yariv reagent (βGlcY). Already in the 1970s, the occurence of AGPs in extracts of different bryophytes and ferns was shown by precipitation in gel-diffusion assays with Yariv’s reagent (Clarke *et al*., 1978). Additionally, AGPs in plant tissue or extracts can be detected using monoclonal antibodies directed against AG motifs. Using these antibodies in microscopy or ELISA revealed AGP glycan epitopes in different bryophytes (Berry *et al*., 2016; Kremer *et al*., 2004; Ligrone *et al*., 2002; Bartels *et al*., 2017; Happ and Classen, 2019) and ferns (Eeckhout *et al*., 2014; Lopez and Renzaglia, 2014; Bartels and Classen, 2017).

Up to now, fine structures of AGPs from bryophytes (Bartels *et al*., 2017; Fu *et al*., 2007, Happ and Classen, 2019) and ferns (Akiyama *et al*., 1987, Bartels and Classen, 2017) were only sparsely investigated. To systematically assess cell wall characteristics of the emerging model ferns, we isolated and characterized AGPs from the leptosporangiate ferns *Azolla filiculoides*, *Salvinia molesta* and *Ceratopteris richardii*. To obtain broader insight into AGP biosynthesis in ferns, we searched for members of the glycosyltransferase family 31 (GT31) involved in AG glycan biosynthesis as well as identified sequences encoding AGP and other HRGP protein backbones. Knowledge of cell wall diversity contributes to our understanding of plant evolution. Due to their crucial phylogenetic position as sister to land plants, ferns are key for understanding cell wall evolution of land plants.

## RESULTS

We used *Salvinia molesta*, *Azolla filiculoides*, and *Ceratopteris richardii* to investigate the diversity and conservation of AGPs in cell walls of ferns. The three species were cultivated under greenhouse conditions, harvested, and dried (see Figure 1 for habit of cultivated ferns and phylogeny of the green plant lineage). In the water soluble fractions (AE) of all species (Tables S1-S3), Gal and Glc were present in high amounts. The content of Ara differed between the investigated species with around 30 % in *Salvinia*, 20 % in *Ceratopteris,* and only 10 % in *Azolla*. AE of *Azolla* further differed by higher amounts of Man, uronic acids and Fuc as well as lack of 3-*O*-methylrhamnose (3-*O*-MeRha; trivial name acofriose). The content of Xyl was another striking difference between the species with low amounts in *Salvinia* but over 20 % in *Ceratopteris* and *Azolla*.

**Figure 1.**
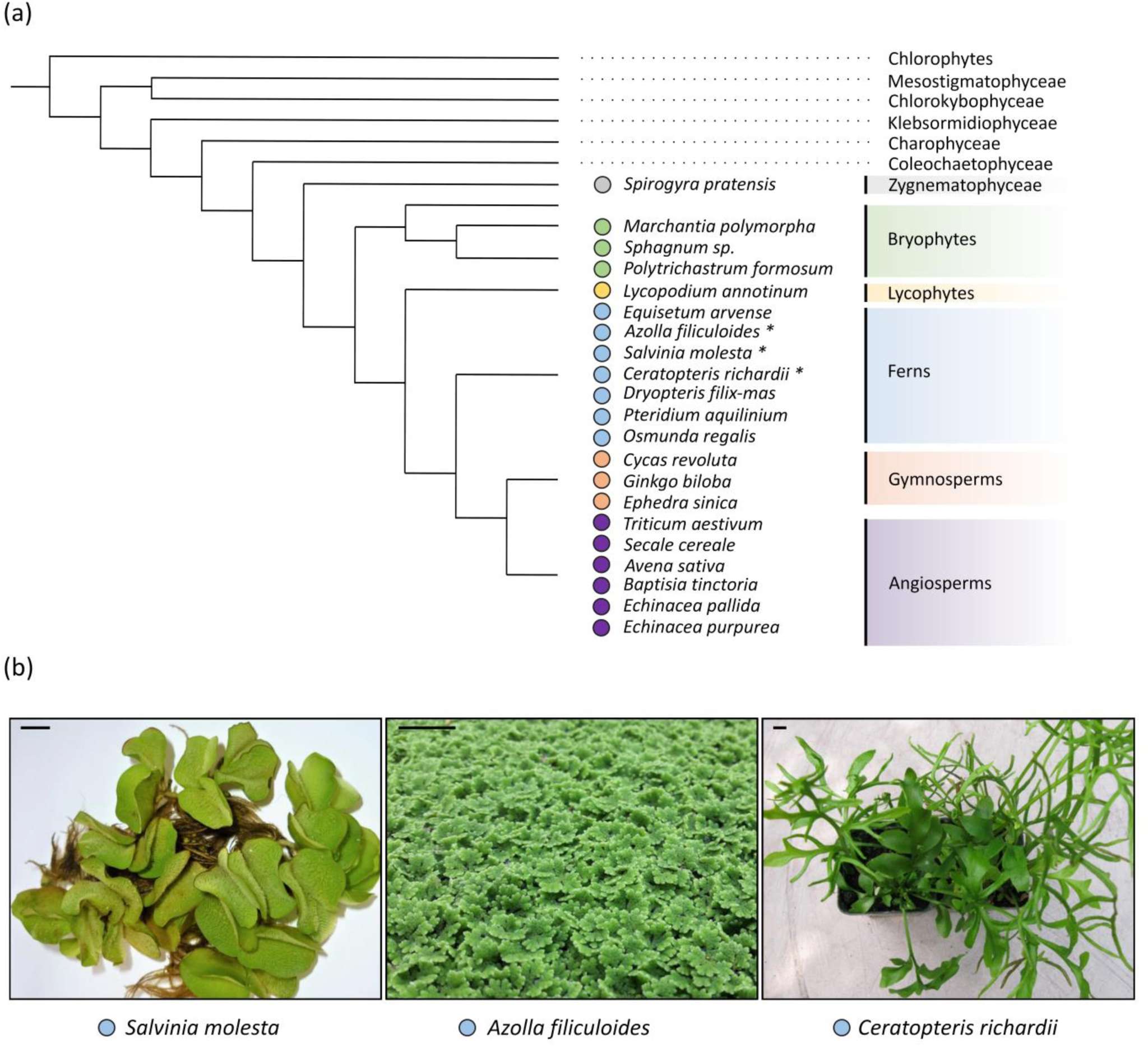
Overview of the investigated plant species. (a) Phylogenetic position of the analyzed plants according to de Vries and Archibald (2018) and Puttick *et al*. (2018). The colored dots indicate inclusion in the PCA (see Figure 3). The fern species which were cultured for this study are marked by asterisks. (b) Cultivation of the ferns *S. molesta*, *A. filliculoides* and *C. richardii* in the greenhouse of the Pharmaceutical Institute of Kiel University. Scale bars = 1 cm.

### (Glyco)diversity in fern AGPs

In a gel diffusion assay with the water-soluble fractions (AE) of the three ferns and βGlcY, red precipitation lines occurred after 22 h of incubation in the dark at room temperature, indicating the presence of AGPs (Figure 2a). Presence of AGPs was further confirmed by isolation with βGlcY in amounts of 0.08 % (*Salvinia*, n=3), 0.18 % (*Azolla*, n=2) and 0.07 % (*Ceratopteris*, n=1) in relation to the dry weight of the ferns.

**Figure 2.**
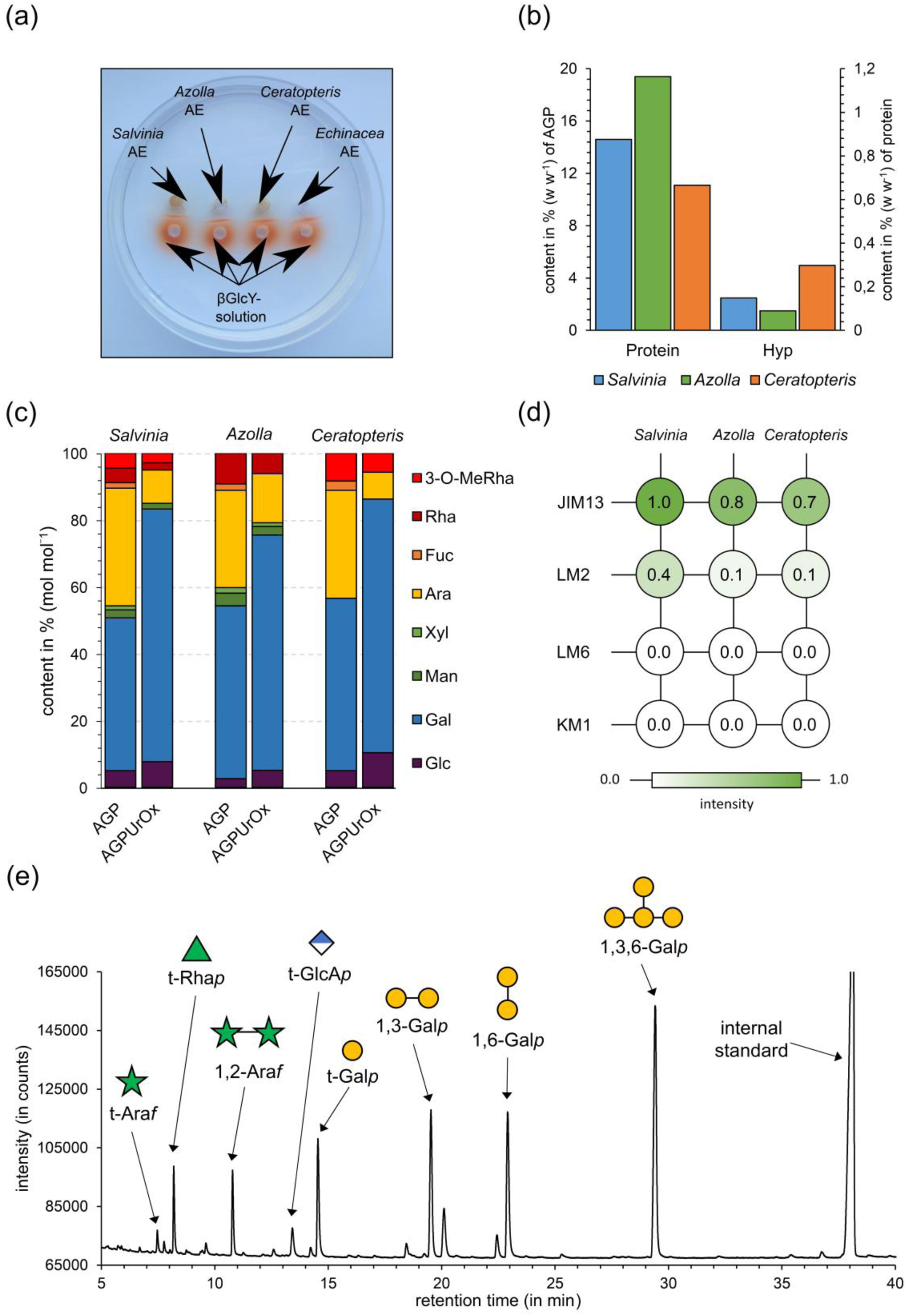
Features of AGPs from *S. molesta*, *A. filiculoides* and *C. richardii*. (a) Gel diffusion assay with βGlcY and fractions AE from the three ferns (100 mg mL^-1^) compared to *Echinacea purpurea* AGP (10 mg mL^-1^). The red precipitation line indicates presence of AGPs. (b) Protein (% w w^-1^ of AGP) and hydroxyprolin content (% w w^-1^ of protein) in AGPs from the three species. (c) Relative monosaccharide composition of AGPs from the three ferns in the native state (AGP) and after reduction of uronic acids and partial hydrolysis with oxalic acid (AGP_UrOx_) determined by gas chromatography (GC) (% mol mol^-1^). (d) Results of ELISA experiments with AGPs and antibodies directed against epitopes present in AGPs (JIM13, LM2, LM6, KM1). For epitopes of the antibodies see Experimental section. (e) FID chromatogram of partially methylated alditol acetates (PMAAs) from *Salvinia* AGP_UrOx_ after GLC analysis.

The amounts of nitrogen in the AGP samples were determined by elemental analyses and used to estimate the protein content of the AGPs according to Kjeldahl (factor 6.25, Figure 2b). The protein amounts of AGPs were 14.6 % for *Salvinia*, 19.4 % for *Azolla,* and 11.1 % for *Ceratopteris*. In AGPs, the amino acid hydroxyproline is responsible for *O*-glycosidic linkage of the arabinogalactan moieties to the protein. Photometric quantification of this amino acid (Stegemann and Stalder, 1967) revealed amounts of 0.36 % for *Salvinia*, 0.29 % for *Azolla*, and 0.55 % for *Ceratopteris* AGP. This means that Hyp accounts for 2.5 % of the protein moiety in *Salvinia*, 1.5 % of the protein moiety in *Azolla*, and 5.0 % of the protein moiety in *Ceratopteris* AGP (Figure 2b).

As expected, Gal and Ara were the dominant monosaccharides of the Yariv-precipitated AGPs and accounted for slightly over 80 % of the neutral monosaccharides in all species (Figure 2c, Table S4) with an Ara : Gal ratio comparable to other AGPs of ferns (Bartels and Classen, 2017) and seed plants (Clarke *et al*., 1978). These monosaccharides were accompanied by Rha / 3-*O*-MeRha in substantial amounts of together between 8.3 % and 9.2 %. Interestingly, the ratio between both deoxyhexoses varied strongly. In *Salvinia* AGP, both were present in nearly equal amounts, whereas in *Azolla* AGP, 3-*O*- MeRha was present only in traces and in *Ceratopteris* AGP, the methylated Rha is strongly dominating (Figure 2c, Table S4). The monosaccharides Glc, Man, Xyl and Fuc were present in smaller amounts, but it cannot be fully excluded that they might be part of other polysaccharides not completely separated during Yariv precipitation. Some Glc might also be the residue of the precipitating agent βGlcY. Photometric quantification of uronic acids revealed a content of 11.0 % for *Salvinia*, 5.6 % for *Azolla* and 11.1 % for *Ceratopteris* AGPs.

To gain further information, uronic acids were carboxy-reduced to the corresponding neutral monosaccharides with sodium borodeuteride. Afterwards, deuterated Glc was detected, thus revealing that GlcA has been part of the native AGP. Furthermore, the samples were partially hydrolysed with oxalic acid, a treatment known to hydrolyse mainly furanosidic arabinose residues.This led to strong decrease of the Ara content (Figure 2c, Table S4) and also to further decrease of the minor monosaccharides Rha, Fuc, Xyl and Man.

### Fern AGPs structural characteristics appear evolutionarily more flexible compared to angiosperms

Linkage types of monosaccharides were determined by methylation analyses before and after partial acid hydrolysis and uronic acid reduction (Table 1, Figure 2e). In the native AGPs, the typical galactan backbone known from seed plant AGPs was present with pyranosidic Gal in 1-, 1,3- 1,6- and 1,3,6- linkage. According to the retention time and the mass spectra, a further peak represented a mixture of pyranosidic, 1,2- and 1,4-linked hexoses. 1,2 and/or 1,4-linked Gal*p* have also been detected in some moss and other fern AGPs (Bartels *et al*., 2017; Bartels and Classen, 2017). Furanosidic Ara, which is typically located at the periphery of the AGP molecule, was 1-, 1,2- and 1,5-linked with dominance of 1,2-linkage type. After partial acid hydrolysis, there was complete loss of 1,5-linked Ara*f*, strong reduction of terminal Ara*f* and moderate reduction of 1,2-Ara*f*. Furthermore, acid hydrolysis strongly increased the amount of 1,6-linked Gal*p*, which is an indication that Ara is mainly bound to Gal at C-3 of 1,3,6-linked Gal. The increase in terminal Gal is a hint that some Ara or acid-labile Rha or Fuc is bound to C-6 of Gal in the native AGP. Comparable to other fern AGPs, Rha*p* (including 3-*O*-Me-Rha*p*) is localized terminally (Bartels and Classen, 2017). After carboxy-reduction of uronic acids to the corresponding deuterium-labeled neutral monosaccharides, GlcA was detected as a terminal monosaccharide (AGP_UrOx_).

**Table 1.** Linkage type analysis of AGPs and hydrolysed AGPs (AGP_UrOx_) from S. molesta, A. filiculoides and C. richardii in % (mol mol^-1^, n=1).

| Mono-saccharide | linkage type | *S. molesta* |  | *A. filiculoides* |  | *C. richardii* |  |
| --- | --- | --- | --- | --- | --- | --- | --- |
|  |  | AGP | AGP<sub>UrOx</sub> | AGP | AGP<sub>UrOx</sub> | AGP | AGP<sub>UrOx</sub> |
| Galp | 1,3,6- | 30.7 | 33.8 | 27.0 | 30.8 | 35.9 | 37.8 |
|  | 1,6- | 2.1 | 15.8 | 2.6 | 11.5 | 2.2 | 12.4 |
|  | 1,3- | 15.2 | 14.4 | 15.5 | 18.7 | 16.4 | 15.0 |
|  | 1- | 2.1 | 12.0 | 3.4 | 5.1 | 6.0 | 9.4 |
| Araf | 1,5- | 3.9 | - | 4.4 | - | 7.3 | - |
|  | 1,2- | 17.2 | 6.7 | 16.9 | 10.7 | 11.5 | 3.9 |
|  | 1- | 9.6 | 1.8 | 8.5 | 2.6 | 6.4 | 1.8 |
| Rhap | 1- | 11.0 | 5.9 | 9.5 | 6.1 | 7.8 | 8.2 |
| Fucp | 1- | - | - | 1.4 | - | 2.2 | - |
| GlcAp | 1- | nd | 3.6 | nd | 2.9 | nd | 3.2 |
| Manp | 1,6- | 2.9 | - | 2.9 | 1.2 | 1.3 | 2.7 |
| Hexp | 1,4-/1,2- | 5.3 | 6.0 | 7.9 | 9.2 | 3.0 | 5.6 |

nd: not detectable due to the method

Antibodies directed against epitopes of the glycan part of angiosperm AGPs (see Experimental section) were tested for their binding capacities to the different fern AGPs to gain further information on AGP structures (Figure 2d). In the tested concentrations, there was no binding of the fern AGPs to KM1 and LM6, thus supporting structural differences to angiosperm AGPs. For KM1 (Classen *et al*., 2004), it has been shown that β-d-1,6-Gal*p* is part of the epitope (Ruprecht *et al*., 2017), which is present in the fern AGPs only in small amounts. LM6 is directed against α-l-1,5-linked Ara*f* (Verhertbruggen *et al*., 2009) present in arabinans but also in some AGPs. Although this Ara linkage type is present in the investigated fern AGPs, the amounts are lower compared to the 1,2-linkage type. Weak binding of LM6 to *Ceratopteris* AGP was observed after longer times of incubation (data not shown), thus confirming higher amounts of 1,5-Ara*f* in *Ceratopteris* compared to *Salvinia* and *Azolla* AGP. LM2 showed moderate binding to the fern AGPs, probably due to presence of terminal GlcA*p*, which has been described as part of the epitope (Ruprecht *et al*., 2017). JIM13 strongly binds to the three fern AGPs, especially to *Salvinia* AGP. JIM13 is known to recognize many AGPs but the exact epitope is under debate. The trisaccharide β-d-GlcA*p*(1→3)-α-d-GalA*p*-(1→2)-α-l-Rha*f* isolated from hydrolysate of gum karaya binds to this antibody (Yates *et al*., 1996). Strong binding of JIM13 to a rhamnogalactan-protein from *Spirogyra* (Pfeifer *et al*., 2022) further suggests that terminal Rha must be an important part of the epitope of JIM13 and is possibly responsible for binding to the investigated fern AGPs. An exemplary structural proposal for AGP of *Azolla* based on the results above is given in Figure S1.

21 different species representing the streptophyte lineage (a streptophyte alga, three bryophytes, a lycophyte, seven ferns, three gymno- and six angiosperms) were compared with regard to AGP linkage types (Table 2). Here, we only used data from our working group to ensure a comparable methodology for isolation and structural characterization (see Figure 3 for references). The comparison of AGP fine structures over the streptophyte lineage by PCA (Figure 3) revealed first hints on lineage-specific structural features of AGPs. The streptophyte alga *Spirogyra pratensis* and the lycophyte L*ycopodium annotinum* were clearly separated from all other species due to high amounts of terminal and unique 1,3- linked Rha*p* in *Spirogyra* AGP and unusual terminal Ara*p* accompanied by 1,3-linked Ara*f* in *Lycopodium* AGP. Yet, a broader sampling is needed to clarify how common these features are in the respective lineages, or whether these are species-specific characteristics. PCA reveals clustering of angiosperm AGPs due to the common structure with a Gal*p* backbone in 1,3,6- and 1,6- linkage decorated with 1,5-linked and terminal Ara*f*. In contrast, gymnosperms linkage types of AGPs seem to be less conserved, although it needs to be considered that only three species from three lineages, *Ginkgo*, *Cycas* and *Ephedra*, and no representatives from conifers were included in the analyses. For fern AGPs a clear separation to angiosperm AGPs is visible in line with their evolutionary distance. There is an overlap of fern AGPs with those of mosses and gymnosperms. A common feature of AGPs from mosses, ferns and gymnosperms is the presence of the unusual monosaccharide 3-*O*-Me-Rha*p* not observed in any angiosperm AGP or any other angiosperm cell wall polysaccharide. Within the ferns, the only eusporangiate fern AGP from *Equisetum* is separated from the leptosporangiate AGPs. The leptosporangiate *Osmunda* AGP is located in between *Equisetum* AGP and the other leptosporangiate AGPs. This is in accordance with the phylogeny of ferns, where *Osmunda* is sister to the rest of the leptosporangiates. *Azolla* and *Salvinia* are members of the same family (Salviniaceae) and the similarities in their AGPs reflect their close phylogenetic relationship or the similarity of their aquatic lifestyle. Indeed, *Pteridium* and *Ceratopteris* belong to the same family (Pteridaceae), yet *Pteridium* AGPs cluster with those from *Dryopteris*, while those of the aquatic fern *Ceratopteris* is more similar to AGPs from *Salvinia* and *Azolla*.

**Figure 3.**
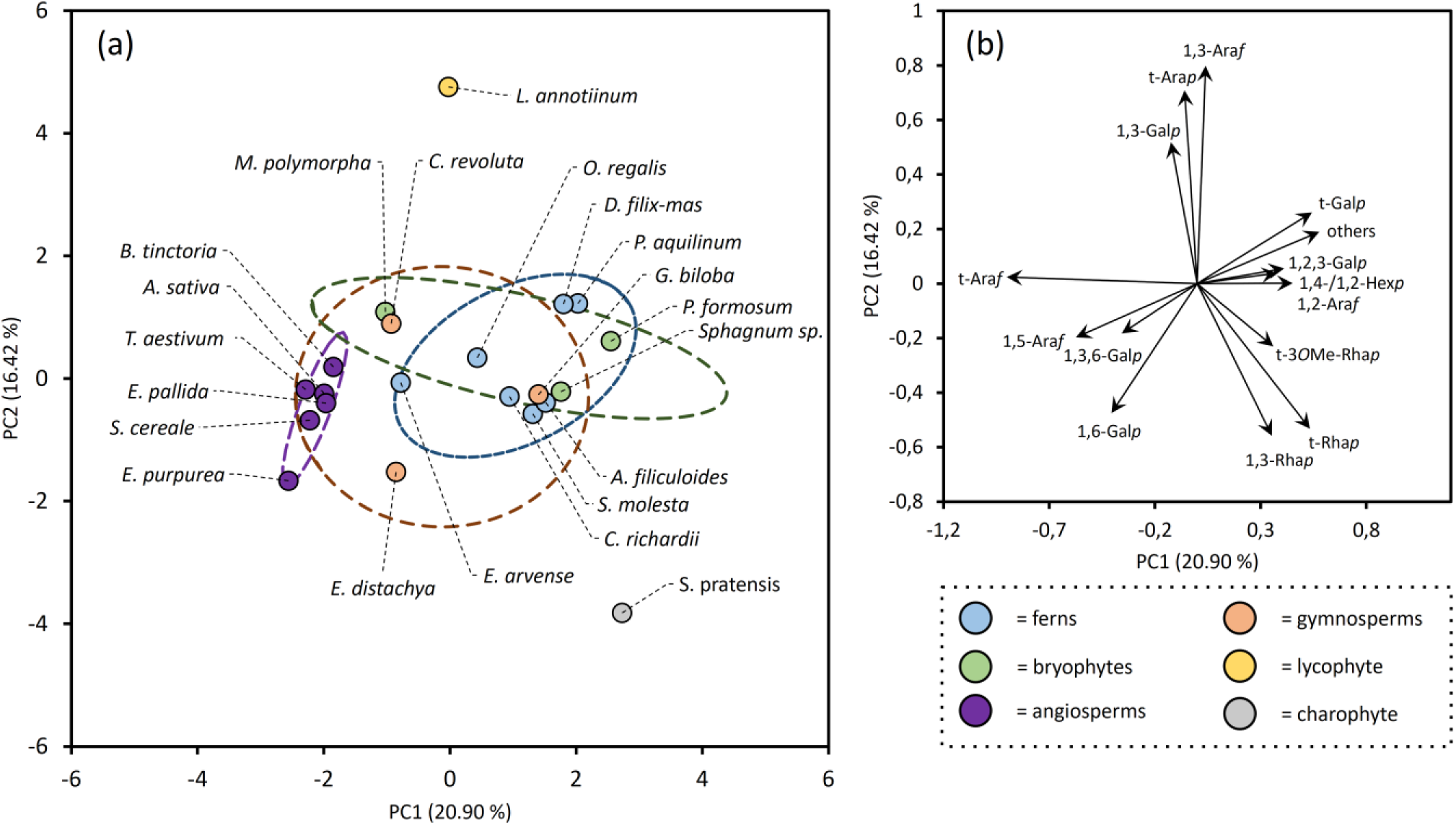
Results of the PCA analysis of linkage-types in AGPs throughout the streptophyte lineage. (a) Scatter plot of PC1 (20.90%) against PC2 (16.42). Different colors were used for each evolutionary group. Ellipses represent 90 % confidence intervals. For full plant names refer to Bartels and Classen (2017), Bartels *et al*., (2017), Baumann *et al*. (2021), Classen *et al*. (2000), Goellner *et al*. (2010, 2011, 2013), Happ and Classen (2019), Pfeifer *et al*. (2022), Thude and Classen (2005) and Wack *et al*. (2005). (b) Loading plot for the PCA with influences of the different input variables.

**Table 2.**
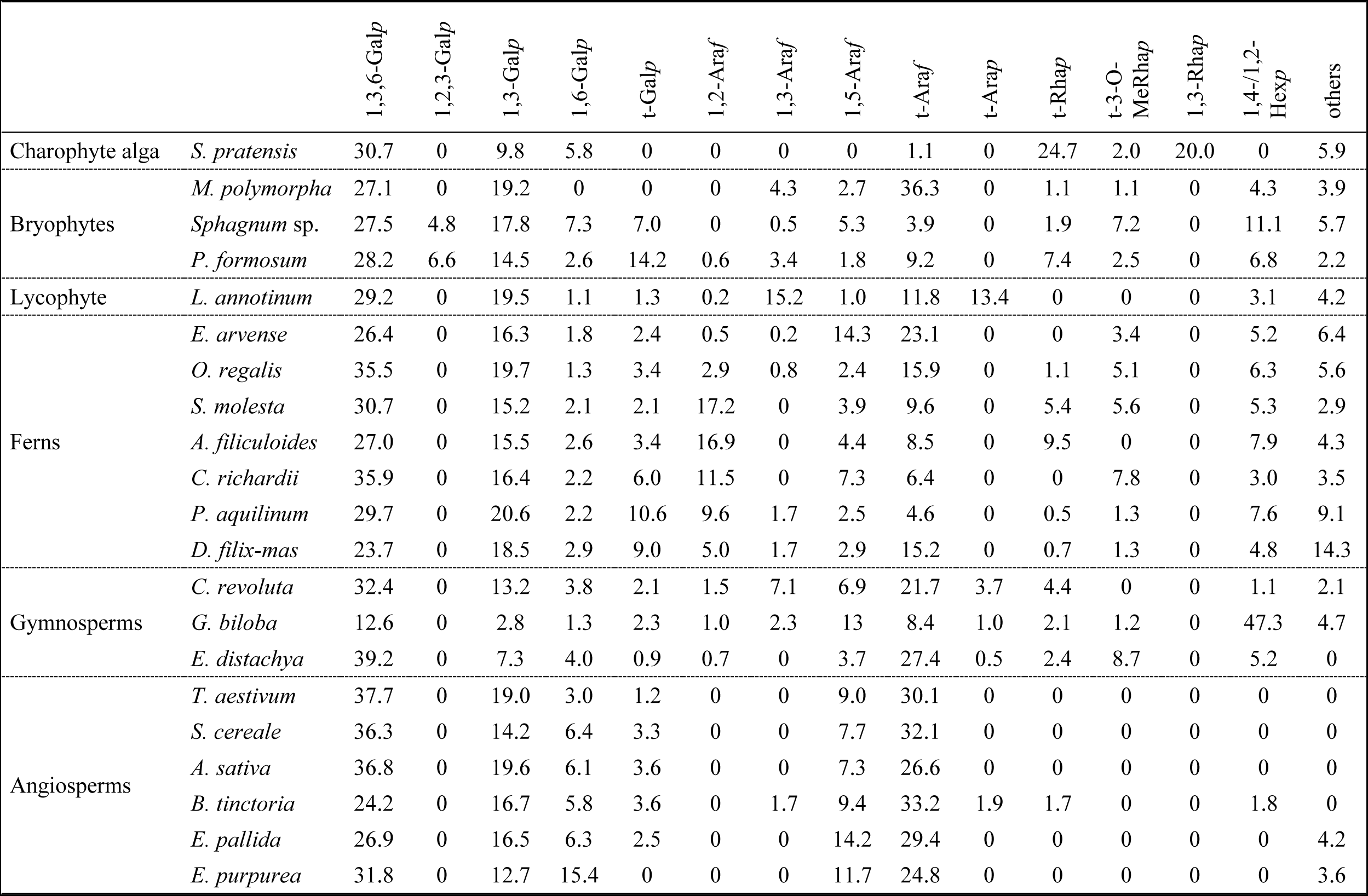
Linkage type analysis of AGPs throughout the streptophyte lineage in % (mol mol^-1^).

### A genetic framework for AGP glycosylation in ferns

Glycosylation of AGPs is very complex and involves addition of large arabinogalactan (AG) moieties to hydroxyproline residues often arranged in characteristic dipeptide repeats: Ala-Hyp, Ser-Hyp, Thr-Hyp. The AGs consist of β-1,3-Gal*p* backbones with β-1,6-linked Gal*p* side chains that are further decorated with mainly Ara but also GlcA, Rha, Fuc or Xyl (for review, see Showalter and Basu, 2016; Silva *et al*., 2020, Strasser *et al*., 2021). The responsible glycosyltransferases (GTs) belong to different GT families that have been classified in the carbohydrate active enzymes (CAZy) database (http://www.cazy.org). Although several GTs responsible for AGP glycosylation have been identified in *Arabidopsis*, many enzymes are still unknown, e.g. most arabinosyl- and all rhamnosyltransferases (Silva *et al*., 2020). Responsible for galactosylation of AGPs are enzymes of the family GT31. A phylogeny of the *Arabidopsis thaliana* sequences in this family divides into three distinct groups which were subdivided into 9 subgroups I-IX (group A with clades I-IV, group B with clades V and VI, group C with clades VII-IX, Qu *et al*., 2008). Eleven of these enzymes belonging to clades II, III, V and VI have already been characterized and are involved in AGP galactosylation in *Arabidopsis thaliana* (Silva *et al*., 2020).

To elucidate whether homologs of these enzymes are present in ferns, *Arabidopsis* sequences from group A and B were used to search for GT31 sequences in the genomes of *Salvinia*, *Azolla* and *Ceratopteris* as well as in transcriptomic data from other ferns (Carpenter *et al*., 2019; Leebens-Mack *et al*., 2019, Figure 4, Table S5 and Figure S2). The first step of AGP galactosylation is the transfer of the first Gal onto Hyp by a hydroxyproline *O*-β-galactosyltransferase. In *Arabidopsis*, GALT 2-6 (Basu *et al*., 2015a; Basu et al, 2015b) belonging to clade VI as well as HPGT1, HPGT2 and HPGT3 from clade III within GT31 (Ogawa-Ohnishi and Matsubayashi, 2015) have been identified to be responsible for this step of the biosynthesis. Whereas GALT2-6 contain a galectin domain, this domain is absent in HPGT1- 3 (Showalter and Basu, 2016). Genomes of *Azolla*, *Salvinia* and *Ceratopteris* and transcriptomes of all ferns contain at least one corresponding sequence in one of these GT31 families, thus supporting that the transfer of Gal onto Hyp is possible in all fern AGPs.

**Figure 4.**
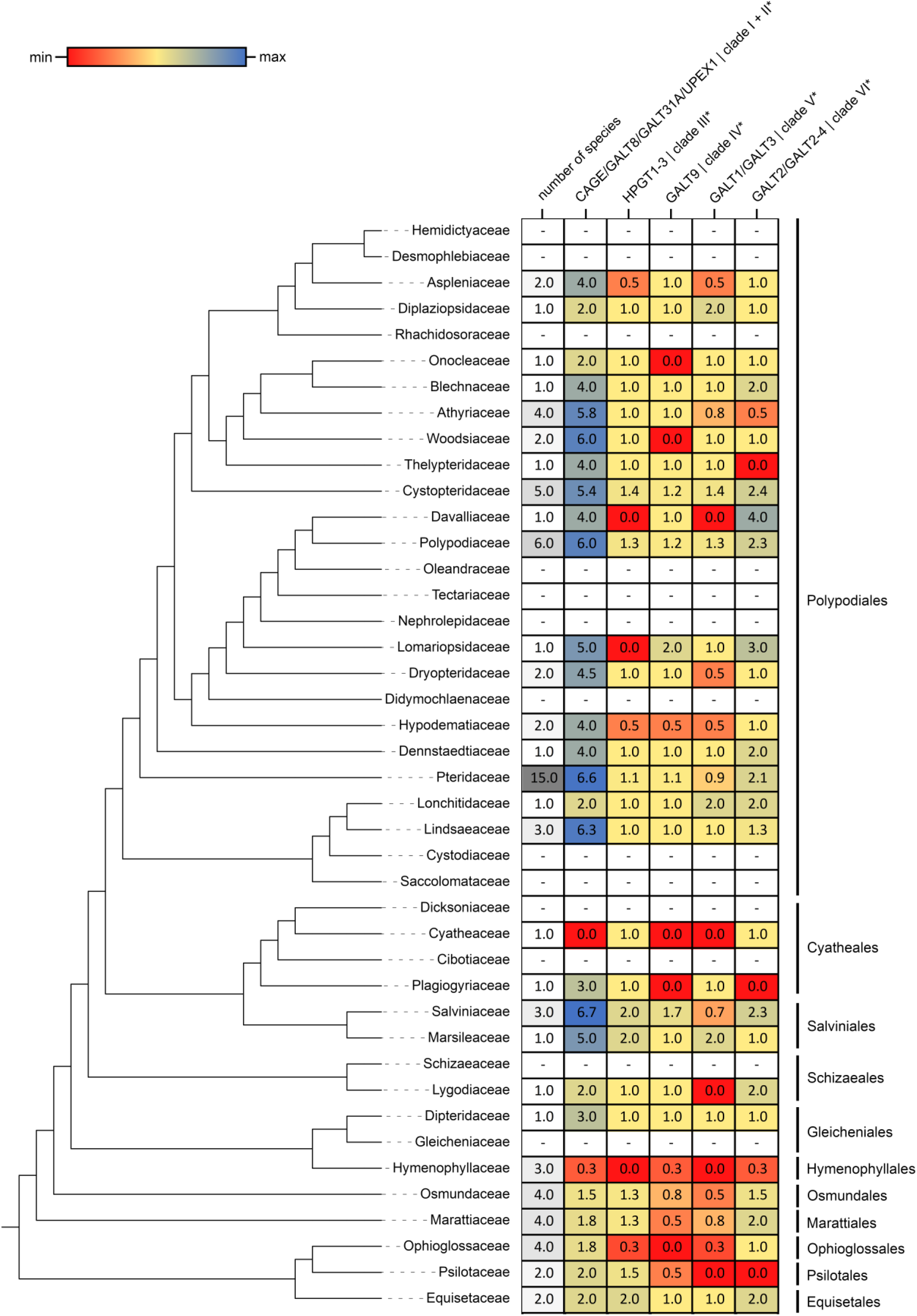
Overview of GT31 homologs from fern transcriptomes and genomes. The fern phylogeny on the left side is based on the dataset of Nitta *et al*. (2022) and was visualized with iTOL (v6.6, Letunic and Bork, 2021). The heatmap on the right side is based on the average homolog number in the respective clades of the phylogenetic tree (Figure S2). White fields with dashes were not covered by the dataset. Clade annotations marked with asterisks are used according to Qu *et al*. (2008).

The *Arabidopsis* genes At1G77810 (Qu *et al*., 2008), At1G33430 (called UPEX1 or KNS4; Suzuki *et al*., 2017) as well as GALT8 (Narciso *et al*., 2021) belonging to clade II and possibly CAGE1 and CAGE2 (clade I, Nibbering *et al*., 2022) encode β-1,3-galactosyltransferases which function in AGP β-1,3-galactan backbone synthesis. Galactan side chains of AGPs consisting of β-1,6-linked Gal are synthesized by β-1,6-galactosyltransferases of family GALT31A (clade II, Geshi *et al*., 2013), although in a recent study GALT31A was found to galactosylate substituted and unsubstituted β-1,3-galactan oligosaccharides rather than β-1,6-linked galactan oligosaccharides (Ruprecht *et al*., 2020). Homologs to sequences from all *Arabidopsis* enzymes from group A and B responsible for synthesis of the galactan structure are present in *Azolla*, *Salvinia*, *Ceratopteris* and all investigated fern transcriptomes. Clades I and II could not be phylogenetically separated (Figure S2; although clade II had also no support as an independent group in Qu et al. 2008, where the groups were originally defined). On balance, the evolution of GT31 group I and II is characterized by lineage-specific diversification (Figure 4): *Arabidopsis* appeared to have slightly more homologs (8 sequences) compared to leptosporangiate (on average 4.8 per species) and eusporangiate ferns (on average 1.8 per species). That said, the investigated fern genomes contain similar numbers of homologs as *Arabidopsis* (8 in *Azolla*, 6 in *Salvinia* and 10 in *Ceratopteris*, see Table S5), indicating that the overall lower number of homologs in transcriptomes of ferns is likely an underestimate caused by technical limitations or other reasons like e.g. low expression or differential regulation. To summarize, the genetic toolkit necessary for galactosylation of Hyp and the biosynthesis of the typical AGP galactan scaffold known from seed plants is present in the fern lineage, corrobarting our analytical findings (Figure S1). Overall, this suggests that Hyp galactosylation and the synthesis of typical AGP galactan is conserved and was already present in the LCA of ferns and seed plants.

### Identification of AGP and other HRGP protein sequences in fern genomes

Sequences for classical AGPs and other HRGPs (extensins and PRPs) as well as hybrid sequences with motifs from different HRGPs were identified (Figure 5) using the established motif and amino acid bias ‘MAAB’ classification system (Johnson *et al*., 2017a,b). Class 1 and 4 comprise classical AGPs with and without GPI anchor respectively, class 2 and 9 are extensins (cross-linking or with GPI-anchor) and class 3 contain PRPs. All other classes consist of hybrid HRGP sequences. For the heterosporous ferns *Azolla* and *Salvinia*, AGP sequences identified were similar in number with 6 for *Azolla* (none with GPI-anchor) and 7 for *Salvinia* (3 with GPI-anchor), whereas for homosporous ferns *Adiantum*, *Alsophila* and *Ceratopteris*, numbers were much higher with 19 - 41 AGP sequences (2 - 7 with GPI-anchor). The same trend was found for extensins with no sequence identified in *Azolla*, only 1 sequence in *Salvinia* and higher numbers of sequences in *Adiantum* and *Ceratopteris*. PRPs were absent in all investigated species and only 1 sequence was detected in the *Ceratopteris* genome. In *Azolla* and *Salvinia* only 1-2 hybrid HRGPs were present, whereas in the homosporous ferns, numerous sequences were found, especially in classes 6 (AGP bias), 10 and 12 (extension bias) and 18 (PRP bias). Sequences belonging to class 24 have been detected in all species with the exception of *Alsophila* but have low percentages of HRGP motifs and possibly comprise likely either non-HRGPs or unknown HRGPs (Johnson *et al*., 2017b)

**Figure 5.**
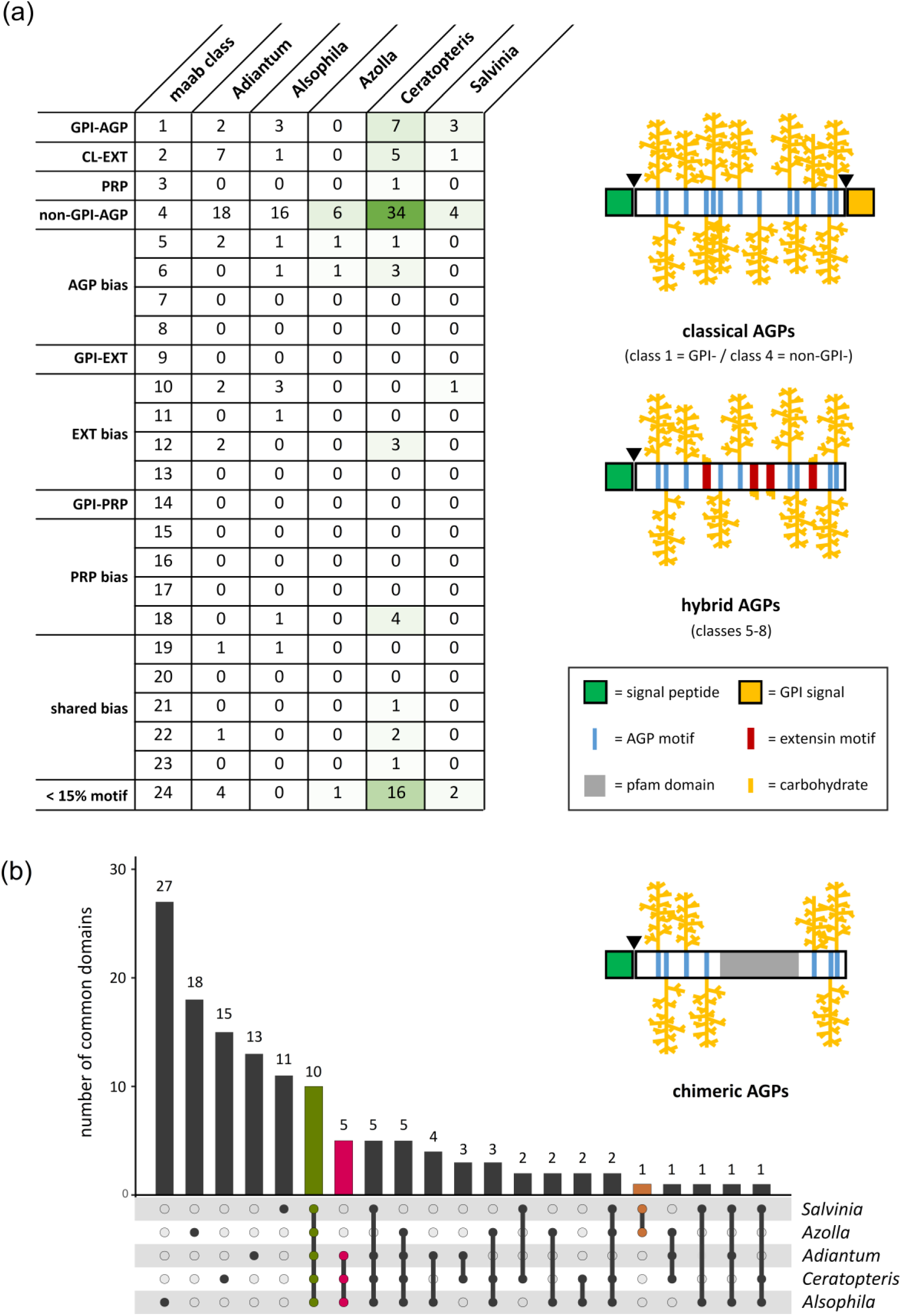
Results of the bioinformatic analyses on AGP backbone sequences from the five fern genomes. (a) MAAB classification of all HRGPs within a possible 24 distinct classes. The schematics on the right side highlight the predicted post-translational modifications on classical, hybrid and chimeric AGPs. (b) Chimeric AGPs are visualized as UpSet plot. Number of common domains of all (green), of all aquatic heterosporous (orange) and all homosporous (magenta) ferns are highlighted. For detailed knowledge of domains, see Figure S3.

Chimeric AGPs are characterized by AGP-motifs, a signal peptide and one or more known functional protein domains. They were identified by screening the translated genome for sequences which meet the criteria (i) signal peptide (SignalP 5.0), (ii) predicted AG regions, and (iii) at least one detectable protein domain (Pfam). 108 chimeric AGPs were present in *Adiantum*, 257 in *Alsophila*, 75 in *Azolla*, 184 in *Ceratopteris*, and 71 in *Salvinia* (Figure S3 and Figure 5b).

By evaluation of the intersections between chimeric AGPs from all five species, shared domains were detected (highlighted in Figure S3 and Figure 5b). 10 domains were present in all ferns. Interestingly, they comprise between 47.1 % and 54.9 % of all chimeric AGPs in ferns (Figure S3). The most dominant members are protein kinase-like AGPs, plastocyanin-like AGPs (Cu bind-like domain, pfam02298) and fasciclin-like AGPs. Another shared group of all ferns are xylogen-like AGPs (containing the LTP-2 domain, pfam14368) which occurs in lower numbers in the small aquatic ferns (each 2 for *Azolla* and *Salvinia*) but higher numbers in *Adiantum* (3 sequences), *Alsophila* (4 sequences) and *Ceratopteris* (7 sequences).

## DISCUSSION

### Polysaccharides of ferns cell walls

Plant terrestrialisation was accompanied by drought stress and leading to diversification of land plants through innovative strategies to reduce dependence on water availability. Structural innovations of tracheophytes include those that enhance transport of water and solutes, as well as taller and turgor- stabilised upright stems for improved spore dispersal and more efficient light capture (Bateman *et al*., 1998; Eeckhout *et al*., 2014). Cell wall modifications are important at all levels of plant growth and development and it is likely that the evolutionary pressures brought about the differences in cell wall composition that occurred during the colonisation of land and the emergence of the tracheophytes. With increasing focus on the molecular processes of plant evolution, different groups have contributed to the knowledge of specific cell wall characteristics in ferns using a variety of methods. Often, immunocytochemical workflows (Eeckhout *et al*., 2014; Leroux *et al*., 2011; Leroux *et al*., 2013b; Leroux *et al*., 2015; Popper 2006; Chernova *et al*., 2020; Harholt *et al*., 2012; Carafa *et al*., (2005)) were applied using various monoclonal antibodies against glycan epitopes (Meikle *et al*., 1994, McCartney *et al*., 2005; Marcus *et al*., 2008; Pedersen *et al*., 2012; Puhlmann *et al*., 1994; Marcus *et al*., 2010). Furthermore, enzyme-assisted digestion with (partial) characterization of the released oligosaccharides was used and shed light on fern xyloglucans (Peña *et al*., 2008; Hsieh and Harris 2012), mannans (Frankova and Fry, 2011; Popper 2006; Silva *et al*., 2011) as well as mixed-linkage glucans (Popper and Fry, 2003; Soerensen *et al*., 2008; Fry *et al*., 2008; Xue and Fry (2012). All these studies fostered the conclusion that fern cell walls share basic features with those of seed plants, such as the ocurrence of cellulose, hemicelluloses and pectic polysaccharides (Leroux *et al*., 2013a; Matsunaga *et al*., 2004; Popper and Fry, 2004; Popper, 2008). Yet they differ from them, for example, in their composition of lignin (Weng *et al*., 2008, see introduction), suggesting that lineage-specific cell wall properties have evolved after the split of the fern and seed plant lineages. These lineage-specific differences seem to occur mainly in the fine-structure of the cell wall, e.g. through variation of side-chain structures. Our focus on AGPs in ferns using methods ranging from carbo-analytical, immunocytochemical to bioinformatic approaches provide detailed insights into the evolutionary history of AGPs in tracheophytes.

### AGPs abound in fern cell walls

Already in the 1970s, the occurence of AGPs in several leptosporangiate ferns was shown by gel- diffusion assay with Yariv’s reagent (Clarke *et al*., 1978). Monoclonal antibodies directed against arabinogalactan epitopes (e.g. JIM4, JIM8, JIM13, KM1, LM2, LM6 or MAC207 (for epitopes and references see Classen *et al*., 2019) were used to verify the presence of AGPs in combination with glycan microarrays, ELISA and microscopy to detect AGPs in different monilophytes (Popper, 2006; Moore *et al*., 2013; Leroux *et al*., 2015; Bartels and Classen, 2017; Eeckhout *et al*., 2014; Lopez and Renzaglia, 2014; Lopez and Renzaglia, 2016). However, reliability of these discoveries are sometimes questionable because the exact structures of epitopes of the antibodies are not always known and cross-reactions with other polysaccharides might occur. For example, LM6 is directed against 1,5-linked Ara*f* and binds to arabinans, yet it can also detect AGPs.

Information on the structure of AGPs from seed plants is extensive, while limited data is available for monilophytes and other seedless plants. Such studies are, to the best of our knowledge, currently limited to seven moss (Geddes and Wilkie, 1971; Kremer *et al*., 2004; Lee *et al*., 2005; Fu *et al*., 2007; Happ and Classen, 2019) and four monilophyte species (Bartels and Classen, 2017). To gain further information on fern AGPs and insight into AGP evolution, we isolated AGPs from *Salvinia*, *Azolla* and *Ceratopteris* by specific precipitation with βGlcY. The amount of *Azolla* AGP was comparable to AGP content of the mosses *Sphagnum* and *Physcomitrium* (Bartels *et al*., 2017) and slightly higher compared to other monilophytes (in mean 0.12 % – 0.15 %, Bartels and Classen, 2017) whereas AGP amounts of *Salvinia* and *Ceratopteris* were slightly lower and comparable to AGP recovery in the moss *Polytrichastrum* (Bartels *et al*., 2017). However, variation in AGP amounts is observed at different times of harvest (Bartels and Classen, 2017); thus it cannot be excluded that some of the observed variation may stem from different times of harvest.

With regard to the AGP protein moieties (high protein and low hydroxyproline content) AGPs from the three fern species investigated here are more similar to some moss AGPs (Bartels *et al*., 2017; Happ and Classen, 2019) than to those from other ferns (Bartels and Classen, 2017). This may reflect similarities in habitat of the mosses and the water ferns we investigated, including characteristics such as high moisture.

The carbohydrate backbones of AGPs mainly consist of linear chains of β1,3-linked Gal*p* attached to Hyp, and branched with β1,6-linked Gal*p* side chains (Ma *et al*., 2018). Our results reveal that AGPs of the three ferns contain this typical galactan core with Gal*p* in 1-, 1,3- 1,6- and 1,3,6-linkage known from seed plants and also from other fern genera (Bartels and Classen, 2017). A proposal for the structure of the carbohydrate moiety of AGP from *Azolla* is shown in Figure S1. The lower amounts of 1,6-linked Gal*p,* indicating a highly branched structure, may be a special feature of fern AGPs. For example in AGP from the angiosperm *Echinacea*, 1,6-linked Gal*p* was present in high amounts (Classen *et al*., 2000). KM1, an antibody directed against 1,6-linked Gal*p* (Ruprecht *et al*., 2017), has been generated against *Echinacea* AGP (Classen *et al*., 2004). This antibody showed no reactivity with the fern AGPs tested here (Figure 2d), thus supporting low amounts of 1,6-linked Galp in the three species.

We observed several differences to seed plants with regard to the monosaccharides present at the periphery of the fern AGPs. The unusual monosaccharide 3-*O*-Me-Rha is present as terminal monosaccharide in *Salvinia*, *Azolla* and *Ceratopteris* AGPs. 3-*O*-Me-Rha has been detected in cell walls of streptophyte algae, bryophytes, lycophytes, ferns and gymnosperms before (Popper and Fry, 2003; Pfeifer *et al*., 2022; Popper, Sadler and Fry, 2004; Matsunaga *et al*., 2004; Anderson and Munro, 1969), but has never been found in angiosperm cell walls. In *Lycopodium* and different ferns, 3-*O*-Me-Rha occurs in rhamnogalacturonan II (Matsunaga *et al*., 2004). Several studies now show that 3-*O*-Me-Rha is also part of many moss, fern and gymnosperm AGPs (Fu *et al*., 2007; Bartels *et al*., 2017; Bartels and Classen, 2017; Happ and Classen, 2019; Baumann *et al*., 2021), but it was not found in *Lycopodium* AGP (Bartels and Classen, 2017).

Furanosidic Ara is present in all typical AGPs in substantial amounts, but linkage types differ. The terminal linkage is the preferred one while Ara*f* in 1,2-, 1,3- and/or 1,5-linkage occurs less often. AGPs from *Salvinia*, *Azolla* and *Ceratopteris* had high and comparable amounts of 1,2 linked Ara*f* to other leptosporangiate fern AGPs (except the early diverging species *Osmunda regalis*, Bartels and Classen, 2017). This linkage type is, however, have not been described for angiosperm AGPs so far. It is noteworthy that linkage types can also differ within one species. In AGPs from the medicinal plant *Echinacea purpurea*, Ara*f* was present in 1,5-linkage, but secreted AGPs from cell cultures of the same species predominantly formed a 1,3-Ara*f* linkage (Classen, 2007). The latter is not limited only to cell culture AGPs but is also the predominant type in AGPs from *Arabidopsis* leaves (Tryfona *et al*., 2012) and *Zostera* (Pfeifer *et al*., 2020).

### Structure of fern AGPs in light of evolution

The comparison of AGP fine structures over the streptophyte lineage by PCA revealed first hints on lineage-specific structural features of AGPs. Generally spoken, angiosperms are most likely more stable regarding the type of side-chain linkages while the other major (land) plant lineages show more flexibility in this direction. The set of angiosperms included in our PCA is composed of members of the monocot order of Poales as well as the two dicot orders Asterales and Fabales. While this dataset does not cover all of angiosperm diversity, the inclusion of monocots and dicots covers the entire evolutionary distance of angiosperms; thus ruling out that the clustering of the PCA can stem from a sampling bias. Clustering of the aquatic ferns *Ceratopteris*, *Salvinia* and *Azolla* seems to correlate with their aquatic habitat. Overall, the data suggest that both, a close phylogenetic relationship and a very divergent lifestyle can drive the evolution of AGP linkage types.

### AGP glycosyltransferases, AGP as well as other HRGP proteins have diversified in ferns

Transfer of the first Gal onto Hyp as well as biosynthesis of the AGP galactan consisting of a β-1,3- linked Gal with β-1,6-linked Gal side chains is performed by members of the glycosyltransferase family 31. We find homologs of *Arabidopsis* Hyp-*O*-galactosyltransferases, β-1,3-galactosyltransferases, and a β-1,6-galactosyltransferase in the genomes of the three investigated ferns and the transcriptoms of other lepto- and eusporangiate ferns. These data agree with our analytical investigations on fern AGP structure and support that the seed plant-typical AGP galactosylation is present in ferns, and was likely present in the LCA of the two lineages.

The occurence of terminal Rha, partially present as 3-*O*-MeRha is the most striking difference of fern to seed plant AGPs. Up to now, the only rhamnosyltransferase characterized with regard to cell wall biosynthesis is involved in pectic rhamnogalacturonan I synthesis (Takenaka *et al*., 2018). Future characterizations of rhamnosyltransferases acting on AGPs will be illuminating.

Protein backbones of classical AGPs consist of an N-terminal signal peptide, a protein sequence rich in Pro, Ala, Ser and Thr (PAST) residues and may have an additional C-terminal GPI anchor sequence (Schultz *et al*., 2000). *In silico* analyses revealed several homologs to classical AGPs, extensins and hybrid HRGPs in ferns such as *Adiantum*, *Alsophila* and *Ceratopteris* in similar amounts as found in seed plants and slightly lower numbers for the ferns *Azolla* and *Salvinia* (Ma *et al*., 2017; Johnson *et al*., 2017a,b). The latter two have small genomes compared to other ferns (Li *et al*., 2018), which could explain our results. In all ferns, the chimeric AGPs comprise mainly protein kinase-like, phytocyanin- like, fasciclin-like and xylogen-like AGPs, which are well described chimeric AGPs in seed plants (Johnson, 2003; Basu *et al*., 2016; Costa *et al*., 2019; He *et al*., 2019; Ma *et al*., 2022; Shafee *et al*., 2020). Conservation of AG-attachment sites close to these protein domains underlines the necessity of glycosylation for the correct function of the respective domains (Dragićević *et al*., 2020). The group of xylogen-like AGPs is of interest with regard to the evolution of vascular tissues (Motose *et al*., 2004; Kobayashi *et al*., 2011). The characteristic pfam domain is among the top 10 protein domains conserved in all investigated fern genomes. However, the domain was less abundant in the genomes of the heterosporous ferns *Azolla* and *Salvinia*. Such universal occurrence across fern species hints towards a fundamental contribution of AGPs to the process of vascular tissue differentiation (see below); lower numbers in some of the water ferns might be connected to their aqueous habitat. While AGP protein sequence numbers of ferns and their galactosylation are similar to that of seed plants, their oligosaccharide structures at the periphery of the AGP molecules differ between ferns and seed plants. This might result in deviating functions and explain lineage-specific differences in cell wall structures of ferns and seed plants.

### On possible functions of AGPs in ferns

Functional roles of AGPs in seed plants amongst others include cell division, growth, programmed cell death, pattern formation, and plant-microbe interactions (for reviews see Seifert and Roberts, 2007; Ellis *et al*., 2010; Ma *et al*., 2018; Hromadová *et al*., 2021). Interaction of GPI-anchored AGPs with plasma membrane-bound transmembrane proteins of the same cell or soluble AGPs with receptors in adjacent cells have been proposed (Ellis *et al*., 2010). Signalling caused by AGPs might also occur by oligosaccharides enzymatically cleaved from the AG glycan moieties or via Ca^2+^-ions (Lamport *et al*., 2014). Knowledge on the function of fern AGPs is limited, but their roles in gamete and embryo development (Lopez and Renzaglia, 2014, 2016) and desiccation tolerance (Moore *et al*., 2013) have been described previously. In *Ceratopteris richardii* eggs and multiflagellated sperm cells (Lopez and Renzaglia, 2014, 2016) are covered by an extraprotoplasmic AGP matrix. Given that seed plant AGPs bind and release Ca^2+^ at the plasmalemma (Lamport and Varnai, 2013), Lopez and Renzaglia (2016) proposed that AGPs from *C. richardii* could facilitate gamete fusion through Ca^2+^ oscillations. These data together with the well-documented role of seed plant AGPs in these processes (for review see Ma *et al*., 2018; Leszczuk *et al*., 2019) suggest that AGPs have likely facilitated reproduction and embryogenesis already in the LCA of vascular plants. Comparison of hydrated and desiccated leaf material of the resurrection fern *Mohria caffrorum* revealed that arabinose-rich polysaccharides (arabinan-rich pectins and AGPs) protect cell walls against desiccation (Moore *et al*., 2013). The Ara-containing polysaccharides act as plasticizers ensuring the maintenance of hydration during drought stress. When the first land plants evolved from an aquatic ancestor, adaptation to water deficiency was a big challenge; the evolution of different kinds of water-transporting mechanisms was likely key for land plants to establish themselves on land and diversify (Woudenberg *et al*., 2022). Involvement of AGPs in water transport has been suggested for many lineages across the land plant tree of life: antibodies directed against AGP epitopes labelled water-conducting cells of different liverworts and mosses (Ligrone *et al*., 2002). Additionally, xylogen-type genes are present in the moss *Physcomitrium* as well as the lycophyte *Selaginella* (Kobayashi *et al*., 2011). The AGP “xylogen” induces xylem differentiation in suspension cultures of the angiosperm *Zinnia* (Motose *et al*., 2004). In *Arabidopsis* AGP epitopes mark the initial cell that gives rise to proto- and metaxylem (Dolan *et al*., 1995). In roots, stems and leaf stalks of the angiosperm *Echinacea purpurea*, labelling of AGPs is most dominant in the area of the xylem (Bossy *et al*., 2009; Goellner *et al*., 2013). The recurrent use of AGPs in water transporting tissues in bryophytes, lycophytes and angiosperms suggest that this function is also conserved in other lineages, including ferns; more data is however needed to support their involvement in development of water-transporting tissues in ferns.

Due to similarities in overall AGP structure, some functions of AGPs might be conserved in bryophytes and tracheophytes. However, differences in fine structure, especially the arabinogalactan moiety, can also cause functional differences. Here we show that *Salvinia*, *Azolla* and *Ceratopteris* reveal high content of Rha and methylated Rha at the periphery of their AGPs. This feature is similar to the structure of AGPs of the bryophytes *Physcomitrium*, *Sphagnum* and *Polytrichastrum* (Fu *et al*., 2007, Bartels *et al*., 2017). The cell walls of the streptophyte algae *Spirogyra* contain rhamnogalactan-proteins that are structurally similar to AGPs with nearly complete replacement of Ara by Rha (Pfeifer *et al*., 2022). Rha and methylated Rha are less polar compared to Ara. This suggests a change from a less hydrophilic AGP surface with Rha and methylated Rha as terminal groups to a more hydrophilic surface dominated by terminal Ara in vascular plants. Given the less hydrophilic features of AGPs in algae, bryophytes and the water ferns it is logical to assume that these AGPs have alternative functions with regard to hydrophobic interactions (Fu *et al*., 2007). Arabinosylation of existing cell wall polymers has been proposed to be an evolutionary strategy to prevent polymer aggregation during water loss (Moore *et al*., 2013). If true, one may assume that the challenges of a life on land have selected for more hydrophilic AGPs with higher amounts of Ara, allowing for dessication tolerance in tracheophytes. The modificatios observed here for the aquatic ferns may be an adaptation to their reversion to aquatic life.

## Conclusion

During plant terrestrialization, plants acquired molecular adaptations to cope with the terrestrial environment. Tracheophyte innovations like xylem and stabilised upright stems needed modifications in cell wall composition, either through elaboration of ancestral polymers or through synthesis of new components. Our investigations show that AGPs, which are important cell wall components with signalling functions, are present in the three fern species investigated and their basic molecular features are similar to those known from seed plants.

Phylogenetic analysis of relevant GT31 enzymes confirmed that AGP protein backbones and homologs of the basic enzymes responsible for biosynthesis of the galactan core of AGPs are present in ferns likely underpinning the conservation of AGP structures in tracheophytes. An unusual attribute of the three fern AGPs was the presence of 1,2-linked Ara*f* in high amounts which has also been detected in other leptosporangiate fern AGPs, but not in AGPs from other taxa. Interestingly, fern AGPs share a special feature with moss and also gymnosperm AGPs, the occurrence of terminal 3-*O*-MeRha not present in angiosperms. Identification of enzymes that synthesize AG glycans in non-flowering plants, especially rhamnosyltransferases, is a challenge for the future.

More phylodiverse fern genomes are needed to identify AGP protein backbones and glycosyltransferases involved in polysaccharide biosynthesis and AGP glycosylation in this lineage of tracheophytes. Future studies should also include detailed structural analyses of AGPs from higher numbers of bryophytes and monilophytes. Generation of AGP mutants, e.g. by CRISPR/Cas9 technique, will help to gather new information on AGP functions in the streptophyte lineage. As AGPs are possibly involved in water balance of plants, they might be interesting targets to modulate desiccation tolerance of plants in times of climate change.

## EXPERIMENTAL PROCEDURES

### Plant material, growth conditions and sampling

The ferns Salvinia × molesta D. S. MITCHELL (Salvinia, S. molesta), Salvinia auriculata AUBL., Salvinia cucullata ROXB., Salvinia natans (L.) ALL., Azolla filiculoides LAM. (Azolla, A. filiculoides) and Ceratopteris richardii BRONGN. (Ceratopteris, C. richardii) were cultivated in the greenhouse in the Garden of the Pharmaceutical Institute of the Christian-Albrechts-University of Kiel between August and October 2018 (Salvinia), September and January 2019/2020 (Azolla) and July and August 2021 (Ceratopteris); see Figure 1. Spores of Ceratopteris RN3 strain were kind gifts of Péter Szövényi (University of Zurich). For spore germination, gametophyte development and sporophyte culture on C-fern medium, the protocol of Plackett et al. (2015) was used. Plant material was cleaned with water and freeze-dried.

### Isolation of arabinogalactan-proteins (AGPs)

The freeze-dried and ground plant material was extracted two times (2 h and 21 h) with 70 % acetone solution (*V/V*) in a ratio of 1:20 (*w/V*) for *A. filiculoides* and *C. richardii* and in a ratio of 1:30 (*w/V*) for *S. molesta* under constant stirring at 4 °C to remove phenolic compounds. Afterwards, the air-dried plant residue was extracted with double-destilled water (ddH_2_O) in a ratio of 1:20 (*w/V*) for *A. filiculoides* and *C. richardii* and in a ratio of 1:30 (*w/V*) for *S. molesta*. After constant stirring for 21 h at 4 °C, the aqueous extract was separated through a tincture press from the remaining plant material.

The aqueous extract was heated in a water bath at 90 – 95 °C for ten minutes to denature proteins. These proteins were removed by centrifugation (4,122 *g*, 20 min, 4 °C), and the extract was concentrated with a rotary evaporator at 40 °C and 0,010 mbar to approximately one tenth of its volume. In order to precipitate polysaccharides and AGPs, the aqueous extract was poured into ethanol at 4 °C in a ratio of 1:4 (*V/V*) and stored overnight at 4 °C. The precipitated material was isolated by centrifugation (19,000 *g*, 4 °C, 20 min) and freeze-dried (AE).

Following the procedure of Classen *et al*. (2005), AGP was isolated from the AE with the β-Glc-Yariv reagent (βGlcY). After dissolving AE in ddH_2_O and βGlcY (1 mg mL^-1^) in an equal volume of sodium chloride solution (0.3 mol L^-1^), the AGP was precipitated overnight at 4 °C after addition of the βGlcY solution. The AGP-βGlcY-complex was separated by centrifugation (19,000 *g*, 4 °C, 20 min) and re- dissolved in ddH_2_O. After heating to 50 °C, sodium hydrosulfite was added until the red color disappeared. To purify the aqueous AGP solution, it was dialyzed against demineralized water for four days at 4 °C (MWCO 12 – 14 kDa) and freezed-dried afterwards.

### Analysis of monosaccharides

For determination of the neutral monosaccharide composition, a modified method of Blakeney *et al*. (1983) was used. To 1 – 10 mg of sample 1.0 mL trifluoracetic acid (TFA, 2 mol L^-1^) and 50 μL internal standard *myo-*inositol (10 g L^-1^) were added, incubated for 1 h at 121 °C, washed three times with 5 mL ddH_2_O and evaporated. The samples were reduced with 200 μL ammonium hydroxide (1 mol L^-1^) and 1.0 mL sodium borohydride in dimethyl sulfoxide (DMSO, 1 g 50mL^-1^). After incubation for 1.5 h at 40 °C the reaction was stopped with 100 μL glacial acetic acid. For the acetylation, 200 μL of 1- methylimidazole and 2.0 mL of acetic anhydride were added, incubated for 20 min at room temperature and stopped by addition of 10 mL ddH_2_O. The samples were acidified with 1.0 mL aqueous sulfuric acid (0.1 mol L^-1^) and liquid-liquid extraction was performed with an equivalent volume of dichloromethane. The identification and quantification of the neutral monosaccharides was performed by gas chromatography (GC) with flame ionization detection (FID) and mass spectrometry detection (MSD): GC + FID: 7890B; Agilent Technologies, USA; MS: 5977B MSD; Agilent Technologies, USA; column: Optima-225; Macherey-Nagel, Germany; 25 m, 250 μm, 0.25 μm; helium flow rate: 1 ml min^-1^; split ratio 30:1. A temperature gradient was used to achieve peak separation (initial temperature 200 °C, subsequent holding time of 3 min; final temperature 243 °C with a gradient of 2 °C min^-1^). 3-O-MeRha was identified by retention time and mass spectrum (see Happ and Classen, 2019).

According to a modified method of Blumenkrantz and Asboe-Hansen (1973), the uronic acid (UA) amount was determined photometrically using a linear calibration generated with a mixture of glucuronic and galacturonic acid (1:1, *w/w*). 200 μL of a 500 μg L^-1^ sample dissolved in sulphuric acid (4 %, *V/V*) was mixed with 1.2 mL of sodium tetraborate in sulphuric acid (75 mmol L^-1^) and incubated at 100 °C in a water bath for 20 min. After cooling for another 10 min on ice, 20 μL of a solution of *meta*-hydroxy- diphenyl (0.15 %, *w/V*) in NaOH (0.5 %, *w/V*) was added to the sample. After incubation for 15 min at room temperature, the absorbance was determined at 525 nm. For each sample, a threefold determination was performed and an additional blank sample, containing NaOH (0.5 %, *w/V)* instead of the color reagent, was substracted.

### Reduction of uronic acids and partial degradation of AGPs

The AGP fraction was first subjected to uronic acid reduction followed by oxalic acid partial hydrolysis. A modified method of Taylor and Conrad (1972) was used to perform a carboxy-reduction of the uronic acids. After dissolving 20 – 30 mg AGP in 20 mL ddH_2_O, 216 mg of *N*-cyclohexyl-*N*’-[2-(*N*- methylmorpholino)-ethyl]-carbodiimide-4-toluolsulfonate was added slowly under constant stirring until it was completely dissolved. An autotitrator adjusted the pH to 4.75 with 0.01 M HCl and maintained it for another 2 h. The uronic acids were then reduced by the dropwise addition of sodium borodeuteride solutions in increasing concentrations (2.0 mL of 1 mol L^-1^; 2.5 mL of 2 mol L^-1^; 2.5 mL of 4 mol L^-1^). To avoid strong foaming during the reaction, one to two drops of 1-octanol were added beforehand. In addition, the pH value was set to 7.00 with 2 M HCl by an autotitrator and also maintained for another 2 h after the reduction agent was fully added. Finally, the pH was adjusted to 6.5 with glacial acetic acid, the solution was dialysed for three days at 4 °C against demineralized water (MWCO 12 – 14 kDa) and freeze-dried.

For the oxalic acid hydrolysis, a modified method of Gleeson and Clarke (1979) was used. 10 – 15 mg of the uronic acid reduced sample was dissolved in 2.0 mL of oxalic acid (12.5 mmol L^-1^) and incubated for 5 h at 100 °C. After cooling to room temperature, the hydrolysate was precipitated in ethanol at a final concentration of 80 % (*V/V*). The precipitate was separated by centrifugation (19,000 *g*, 10 min, 4 °C), washed twice with 80 % ethanol (*V/V*), dissolved in 2 mL ddH_2_O and freeze-dried.

### Structural characterization of arabinogalactan moiety of the AGP

Structure elucidation of AGPs was performed by a modified method according to Harris *et al*. (1984) with potassium methylsulfinyl carbanion (KCA) and iodomethane (IM) in DMSO. The sample (1-5 mg freeze-dried AGP) was treated stepwise with the KCA and IM (1. 100 µL KCA for 10 min; 2. 80 µL IM for 5 min; 3. 200 µL KCA for 30 min; 4. 150 µL IM for 30 min; 5. 500 µL KCA for 30 min; 6. 400 µL IM for 60 min). The permethylated sample was dissolved in 3.0 mL dichloromethane/methanol in a ratio of 2:1 (*V*/*V*), washed three times with 2.0 mL water and centrifuged (5 min, 2,500 *g*). 2.0 mL 2-,2- dimethoxy propane and 20 µL glacial acid were added to the sample, heated to 90 °C and dried under a stream of nitrogen. The samples werde hydrolyzed with trifluoroacetic acid (2.0 mol L^-1^) at 121 °C for 1 h, mixed with 2.0 mL water and freeze-dried. Afterwards, the partially methylated monosaccharides were reduced by addition of sodium-borohydride (0.5 mol L^-1^), dissolved in ammonium hydroxide (2.0 mol L^-1^), incubated at 60 °C for 1 h, and the reaction stopped by 0.5 mL acetone. Acetylation was performed by addition of 200 µL glacial acid, 1.0 mL ethyl acetate, 3.0 mL acetic anhydride and 100 µL perchloric acid and stopped by demineralized water and 100 µL 1-methylimidazole. The permethylated alditol acetates were extracted with dichloromethane and separated and detected by GLC-mass spectroscopy as described above (instrumentation see above:”Analysis of monosaccharides”; column: Optima-1701, 25 m, 250 µm, 0.25 µm; helium flow rate: 1 mL min^-1^; initial temperature: 170 °C; hold time 2 min; rate 1 °C min^-1^ until 210 °C was reached; rate: 30 °C min^-1^ until 250 °C was reached; final hold time 10 min).

### Principal Component Analysis (PCA)

Based on the resulting biochemical data of the methylation analyses, a PCA was performed with the mean values after standardization by using SigmaPlot (build 14.5.0.101) with default settings. The proportion of the terminal rhamnose (unmethylated) to the terminal 3-*O-*Me-Rha was calculated from the compositional data of the acetylation analyses. See Figure 1 for full plant names and phylogenetic position as well as Figure 3 for references.

### Determination of the protein moiety and the hydroxyproline content

According to the method of Kjeldahl (1883), the protein content was calculated from the nitrogen content (factor 6.25). The nitrogen content was measured by elemental analysis in the Chemistry Department of CAU Kiel University, Kiel, Germany (HEKAtech CHNS Analyzer).

For the quantification of hydroxyproline (Hyp), the methodology of Stegemann and Stalder, 1967 was used with slight modifications. AGPs (10 mg mL^-1^) were hydrolyzed in a 6 M HCl solution at 110 °C for 22 h. After cooling to room temperature, the hydrolysate was centrifuged at 25,000 *g* for 10 min. 160 µL of the supernatant were diluted with 3840 µL ddH_2_O and 0.6 mL of this dilution was measured in triplicate, as well as a blank value. The blank value was mixed with 0.3 mL of oxidation buffer (pH 6.8; 2.6 g citric acid monohydrate; 7.8 g sodium acetate anhydrous; 1.4 g sodium hydroxide; 25 mL 1- propanol filled to 100 mL ddH_2_O) and the samples with 0.3 mL of an oxidating solution (210 mg chloramine T in 15 mL oxidation buffer pH 6.8) and incubated for 20 min at room temperature in the dark. After addition of 0.3 mL colour reagent (1.5 g 4-dimethylaminobenzaldehyde; 5.25 mL perchloric acid, 60 % (*V/V*); 9.25 mL 2-propanol) to all samples, they were incubated for 15 min in a water bath at 60 °C. After cooling under running tap water for 3 minutes, incubation followed for 30 min at room temperature, before the absorbance was measured at 558 nm with an UV/Vis-spectrophotometer. To determine the hydroxyproline content, a linear regression analysis was performed, using 4-hydroxy-l- proline as a standard.

### Indirect Enzyme-linked immunosorbent assay (ELISA)

96-well plates were coated with 100 µl per well of the sample in the concentration 25 µg mL^-1^ in ddH2O and incubated at 37.5 °C for 3 days with open cover. After washing the plates three times with 100 µl phosphate buffered saline (PBS)-T (pH 7.4, 0.05 % Tween® 20) per well, they were blocked with 200 µl of BSA (bovine serum albumin, 1 % (*w/V*)) in PBS per well and incubated again at 37.5 °C for 1 h. The plates were again washed three times with 100 µl of PBS-T per well. After addition of 100 µl primary antibody solution (JIM13, KM1, LM2, LM6, LM10, LM15, LM19, LM20, LM25) per well in 1:20 (*V/V*) PBS 7.4 dilution, the plates were incubated for 1 h at 37.5 °C and washed again three times with 100 µl PBS-T per well. This was repeated with the secondary antibody (anti-mouse-IgG (KM1) or anti-rat-IgG (others) conjugated with alkaline phosphatase, produced in goat, Sigma-Aldrich Chemie GmbH, Taufkirchen, Germany) in a dilution of 1:500 (*V/V*) in PBS 7.4. Finally, 100 µl of the substrate *p*-nitro- phenylphosphate was added per well and the plates were incubated in the dark at room temperature. The absorbance was measured at 405 nm with a plate reader after ten minutes. The samples were analyzed in triplicate. For visualization, the absorbance of the control was set 0 and the highest signal in the dataset was set 1.0. Epitopes of the antibodies and key references are listed in Table S6.

### Gel diffusion assay

In an agarose gel (Tris-HCl, 10 mmol L^-1^; CaCl_2_, 1 mmol L^-1^; NaCl, 0.9 % *w/V*; agarose, 1 % *w/V*), several cavities were stamped. The cavities in the first row were filled with dilutions (100 mg mL^-1^) of each sample and one of them with the positive control, AGP from *Echinacea purpurea* (10 mg mL^-1^). In the second line, the cavities were filled with the βGlcY solution (1 mg mL^-1^). After incubation in the dark at room temperature for 22 h, a red precipitation line appears if the sample contains AGPs.

### Phylogenetic analysis of glycosyltransferase family 31 (GT31)

Putative GT31 homologs of ferns were identified by using the BLASTp interface (E-value of 0.01) at the 1KP database (Carpenter *et al*., 2019; Leebens-Mack *et al*., 2019) webserver as well as blasting the translated genomes of *A. filiculoides* (v1.1), *C. richardii* (v2.1) and *S. cucullata* (v1.2) (Table S7). AGP- relevant GT31 protein sequences from group A and B of *A. thaliana* were used as query sequences against all accessible leptosporangiate and eusporangiate ferns (accessed at June, 23^rd^, 2022). The resulting candidate sequences as well as the query sequences were aligned with L-INS-I using MAFFT (version 7.490; Katoh and Standley, 2013). All sequences with a minimum of 200aa were included in the dataset. the phylogenetic tree was constructed by using IQ-Tree using model JTT+I+G4 identified by model finder in-built in IQ-TREE (multicore version 1.5.5 ; Letunic and Bork, 2021). 100 bootstrap replicates were performed.

### AGP backbone sequence identification and classification

The annotated protein sequences from the ferns *A. filiculoides* (v1.1), *Adiantum capillus-veneris*, *Alsophila spinulosa* (version 5), *C. richardii* (v2.1), *S. cucullata* (v1.2) were downloaded from the respective web resources (Table S7) and imported into the R environment. By using the R-package “ragp” (Dragićević *et al*., 2020), the N-terminal signal sequences were identified and the maab-pipeline (Johnson *et al*., 2017a) as implemented in the R-package was used to classify the proteins based on their motifs and amino acid biases. All small (< 90 amino acids), highly similar (≥ 0.95 similarity) and chimeric sequences (containing any Pfam domain, E-value < 1e-5) were excluded from the classification. To differentiate the classes 1 and 4 the bigPI algorithm (Eisenhaber *et al*., 2003) was used to predict GPI-anchor signals.

Chimeric AGP sequences were identified by detecting (STAGV)P by using the implementation in the “ragp”-package with the strict mode (at least 4 dipeptided in a maximal distance of 4 amino acids) used by Paunović *et al*. (2021). All dipeptides in an annotated pfam domain (by querying the Pfam database *via* the CDD webtool, Lu *et al*., 2020) or in the signal sequence were not count. The candidate sequences were length-filtered (≥ 90 amino acids) and highly-similar sequences (> 0.95 similarity) were discarded. For each translated genome, the chimeric AGPs were grouped by their largest domain and the number of sequences was plotted as bars in Figure S3. An analysis for common domains in the different ferns was performed and visualized by an UpSet plot (Lex *et al*., 2014) with the R-package “UpSetR” (Conway *et al*., 2017).

## ACKNOWLEDGEMENTS

Funding: LP and BC (project-number 440046237; CL448/3-1), JdV (project-number 440231723; VR132/4-1), and PSZ (project-number 440370236; PSLJ1111/1) are grateful for funding within the framework of MAdLand (http://madland.science), priority programme 2237 of the German Research Foundation (DFG). JdV further thanks the European Research Council for funding under the European Union’s Horizon 2020 research and innovation programme (Grant Agreement No. 852725; ERC-StG “TerreStriAL”). Additional funding was received from the Swiss National Science Foundation (grant nos. 160004, 184826, and 212509 to PS); project funding through the University Research Priority Program “Evolution in Action” of the University of Zurich to PS; a Georges and Antoine Claraz Foundation grant to PS.

## AUTHORS CONTRIBUTIONS

BC and LP planned and designed the research. KM, LP and LS cultivated the plant material, in case of *Ceratopteris* from spores delivered by PS. KM, LP and LS performed the extractions as well as carbohydrate and ELISA experiments. LP, SdV, JdV and KJ performed bioinformatic searches. LP created all main text figures. All authors analysed the data (BC, KM, LS and LP: carbohydrate analysis; LP, SdV and JdV: bioinformatics search for glycosyltransferases; LP and KJ: bioinformatics search for AGP protein backbones). BC wrote the draft manuscript with help of KM and LP; all authors revised the manuscript, read and approved the final manuscript.

## DATA AVAILABILITY STATEMENT

All relevant experimental data can be found within the manuscript and its supporting materials. Datasets for bioinformatic analyses can be found here: https://figshare.com/s/50d87902e1fba3ad98c2

## SUPPORTING INFORMATION

Additional Supporting Information can be found in the online version of this article.

**Table S1.** Yields of water-soluble polysaccharides (AE) from *S. molesta*, *A. filiculoides*, and *C. richardii* in % of dry plant material (w w^-1^).

**Table S2.** Neutral monosaccharide composition of water-soluble polysaccharides from *A. filiculoides*, *S*. *molesta* and *C. richardii* in % (mol mol^-1^).

**Table S3.** Colorimetric determination of the content of uronic acids in the water-soluble polysaccharides from *S. molesta*, *A. filiculoides* and *C. richardii* in % (w w^-1^).

**Table S4.** Neutral monosaccharide composition of AGPs and partially hydrolysed AGPs_UrOx_ from *S. molesta*, *A. filiculoides* and *C. richardii* in % (mol mol^-1^).

**Table S5.** Galactosyltransferase sequences of family GT31 present in genomes (*Azolla*, *Salvinia*, *Ceratopteris*) compared to transcriptomes of other ferns.

**Table S6.** Antibodies tested for binding to fern cell wall AGPs.

**Table S7.** Accessed resources for the analysis of translated fern genomes.

**Figure S1.** Proposed structure for the carbohydrate moiety of AGPs from *Azolla filiculoides*. The proposal was derived from the compositional, immunocytochemical, and linkage-type analyses of AGP and hydrolysed AGPs.

**Figure S2.** Phylogeny of GT31 enzymes throughout the different groups of ferns.

**Figure S3:** Chimeric AGPs of all investigated fern genomes ranked by decreasing numbers.

