## Supplementary material for "Fern cell walls and the evolution of arabinogalactan-proteins in streptophytes": Table S1, Table S2, Table S3, Table S4, Table S5, Table S6, Table S7, Figure S1, Figure S2, Figure S3

**Table S1.** Yields of water-soluble polysaccharides (AE) from *S. molesta*, *A. filiculoides*, and *C. richardii* in % of dry plant material (w $w^{-1}$).

| Yields | *S. molesta* | *A. filiculoides* | *C. richardii* |
|---|---|---|---|
| AE | 4.5 | 5.6 | 3.4 |

**Table S2.** Neutral monosaccharide composition of water-soluble polysaccharides (AE) from *A. filiculoides*, *S. molesta* and *C. richardii* in % (mol $mol^{-1}$).

| Neutral monosaccharide | *S. molesta* AE (n=9) | *A. filiculoides* AE (n=3) | *C. richardii* AE (n=3) |
|---|---|---|---|
| 3-*O*-Me-Rha | 2.6 ± 0.2 | tr | 3.0 ± 0.3 |
| Rha | 4.2 ± 0.3 | 5.1 ± 0.1 | 1.9 ± 0.1 |
| Fuc | 2.2 ± 0.2 | 6.5 ± 0.1 | 2.9 ± 0.1 |
| Rib | 2.6 ± 1.0 | tr | 1.2 ± 0.2 |
| Ara | 30.5 ± 2.8 | 10.8 ± 0.6 | 19.8 ± 0.6 |
| Xyl | 5.4 ± 1.1 | 23.5 ± 1.5 | 22.6 ± 0.8 |
| Man | 6.8 ± 0.5 | 12.3 ± 1.0 | 9.0 ± 1.5 |
| Gal | 25.8 ± 1.8 | 21.7 ± 0.3 | 23.7 ± 1.5 |
| Glc | 19.9 ± 1.8 | 20.1 ± 1.1 | 15.9 ± 1.9 |

tr: trace value < 1 %

**Table S3.** Colorimetric determination of the content of uronic acids in the water-soluble polysaccharides (AE) from *S. molesta*, *A. filiculoides* and *C. richardii* in % (w $w^{-1}$).

| Uronic acids | *S. molesta* | *A. filiculoides* | *C. richardii* |
|---|---|---|---|
| AE | 1.8 ± 0.2 | 8.4 ± 0.3 | 4.4 ± 0.1 |

**Table S4.** Neutral monosaccharide composition of AGPs and partially hydrolysed $AGPs_{UrOx}$ from *S. molesta*, *A. filiculoides* and *C. richardii* in % (mol $mol^{-1}$).

| Neutral monosaccharide | *S. molesta* | | *A. filiculoides* | | *C. richardii* | |
|---|---|---|---|---|---|---|
| | AGP (n=5) | $AGP_{UrOx}$ (n=1) | AGP (n=3) | $AGP_{UrOx}$ (n=1) | AGP (n=3) | $AGP_{UrOx}$ (n=1) |
| 3-*O*-Me-Rha | 4.5 ± 0.3 | 2.9 | tr | tr | 8.3 ± 0.7 | 5.7 |
| Rha | 4.3 ± 0.1 | 2.1 | 9.2 ± 0.2 | 6.1 | tr | tr |
| Fuc | 1.7 ± 0.2 | tr | 1.9 ± 0.0 | tr | 2.8 ± 0.2 | tr |
| Ara | 35.2 ± 2.1 | 10.0 | 29.1 ± 0.5 | 14.7 | 32.3 ± 1.0 | 8.0 |
| Xyl | 1.2 ± 0.2 | tr | 1.7 ± 0.1 | 1.1 | tr | - |
| Man | 2.4 ± 0.4 | 1.7 | 3.8 ± 0.1 | 2.6 | tr | tr |
| Gal | 45.7 ± 2.6 | 75.6 | 51.7 ± 0.5 | 70.4 | 51.6 ± 1.7 | 75.9 |
| Glc | 5.0 ± 0.5 | 7.7 | 2.6 ± 0.6 | 5.1 | 5.0 ± 0.3 | 10.4 |
| Ara : Gal | 1 : 1.3 ± 0.2 | 1 : 7.6 | 1 : 1.8 ± 0.0 | 1 : 4.8 | 1 : 1.6 ± 0.1 | 1 : 9.5 |

tr: trace value < 1 %

**Table S5.** Galactosyltransferase sequences of family GT31 present in genomes (*Azolla*, *Salvinia*, *Ceratopteris*) compared to transcriptomes of other ferns

| | | | Clade I + II* | Clade III* | Clade IV* | Clade V* | Clade VI* |
|---|---|---|---|---|---|---|---|
| | | Number of species included | CAGE GALT8 GALT31A UPEX1 | HPGT 1-3 | GALT9 | GALT1 GALT3 | GALT2 GALT4-6 |
| **Genomes** | | | | | | | |
| | *Arabidopsis thaliana* | 1 | 8 | 3 | 3 | 2 | 4 |
| | *Azolla filiculoides* | 1 | 8 | 3 | 2 | 1 | 3 |
| | *Salvinia cucullata* | 1 | 6 | 1 | 2 | 1 | 2 |
| | *Ceratopteris richardii* | 1 | 10 | 2 | 1 | 1 | 3 |
| **Transcriptomes** | | | | | | | |
| **Leptosporangiates** | | | | | | | |
| Polypodiales | Aspleniaceae | 2 | 8 | 1 | 2 | 1 | 2 |
| | Athyriaceae | 4 | 23 | 4 | 4 | 3 | 2 |
| | Blechnaceae | 1 | 4 | 1 | 1 | 1 | 2 |
| | Cystopteridaceae | 5 | 27 | 7 | 6 | 7 | 12 |
| | Davalliaceae | 1 | 4 | 0 | 1 | 0 | 4 |
| | Dennstaedtiaceae | 1 | 4 | 1 | 1 | 1 | 2 |
| | Diplaziopsidaceae | 1 | 2 | 1 | 1 | 2 | 1 |
| | Dryopteridaceae | 1 | 6 | 1 | 1 | 1 | 2 |
| | Elaphoglossaceae | 1 | 3 | 1 | 1 | 0 | 0 |
| | Hypodematiaceae | 2 | 8 | 1 | 1 | 1 | 2 |
| | Lindsaeaceae | 3 | 19 | 3 | 3 | 3 | 4 |
| | Lomariopsidaceae | 1 | 5 | 0 | 2 | 1 | 3 |
| | Lonchitidaceae | 1 | 2 | 1 | 1 | 2 | 2 |
| | Onocleaceae | 1 | 2 | 1 | 0 | 1 | 1 |
| | Polypodiaceae | 6 | 36 | 8 | 7 | 8 | 14 |
| | Pteridaceae | 14 | 89 | 15 | 15 | 12 | 29 |
| | Thelypteridaceae | 1 | 4 | 1 | 1 | 1 | 0 |
| | Woodsiaceae | 2 | 12 | 2 | 0 | 2 | 2 |
| Cyatheales | Culcitaceae | 1 | 3 | 0 | 0 | 2 | 1 |
| | Cyatheaceae | 1 | 0 | 1 | 0 | 0 | 1 |
| | Thyrsopteridaceae | 1 | 4 | 1 | 0 | 2 | 1 |
| Plagiogyriales | Plagiogyriaceae | 1 | 3 | 1 | 0 | 1 | 0 |
| Salviniales | Marsileaceae | 1 | 5 | 2 | 1 | 2 | 1 |
| | Salviniaceae | 1 | 6 | 2 | 1 | 0 | 2 |

| | | | | | | | |
|---|---|---|---|---|---|---|---|
| Schizeales | Anemiaceae | 1 | 3 | 0 | 1 | 1 | 2 |
| | Lygodiaceae | 1 | 2 | 1 | 1 | 0 | 2 |
| Gleicheniales | Dipteridaceae | 1 | 3 | 1 | 1 | 1 | 1 |
| Hymenophyllales | Hymenophyllaceae | 3 | 1 | 0 | 1 | 0 | 1 |
| Osmundales | Osmundaceae | 4 | 6 | 5 | 3 | 2 | 6 |
| **Eusporangiates** | | | | | | | |
| Marattiales | Marattiaceae | 4 | 7 | 5 | 2 | 3 | 8 |
| Equisetales | Equisetaceae | 2 | 4 | 4 | 2 | 2 | 4 |
| Psilotales | Psilotaceae | 2 | 4 | 3 | 1 | 0 | 0 |
| Ophioglossales | Ophioglossaceae | 4 | 7 | 1 | 0 | 1 | 4 |

* according to Qu et al. (2008)

**Table S6.** Antibodies tested for binding to fern cell wall AGPs

| Antibody | Epitope | Key References |
|---|---|---|
| JIM13 | AGP glycan,<br>e.g. β-D-GlcA*p*-(1→3)-α-D-GalA*p*-(1→2)-α-L-Rha | Pfeifer *et al.* (2022);<br>Yates *et al.* (1996) |
| KM1 | (1→6)-β-D-Gal*p* units in AGs type II | Classen *et al.* (2004);<br>Ruprecht *et al.* (2017) |
| LM2 | (1→6)-β-D-Gal*p* units with terminal ß-D-GlcA*p* in AGP | Ruprecht *et al.* (2017);<br>Smallwood *et al.* (1996) |
| LM6 | (1→5)-α-l-Araf oligomers in arabinan or AGP | Verhertbruggen *et al.* (2009) |

**Table S7.** Accessed resources for the analysis of translated fern genomes.

| Plant species | Genome version | Resource |
|---|---|---|
| *Adiantum capillus-veneris* | version 1<br>(accessed October 6th, 2022) | https://figshare.com/s/47be9fe90124b22d3c0e |
| *Alsophila spinulosa* | version 5 | https://figshare.com/articles/dataset/A_spinulosa_genome_rar/19075346/5 |
| *Azolla filiculoides* | version 1.1 | https://fernbase.org/ |
| *Ceratopteris richardii* | version 2.1 | https://phytozome-next.jgi.doe.gov/ |
| *Salvinia cucullata* | version 1.2 | https://fernbase.org/ |

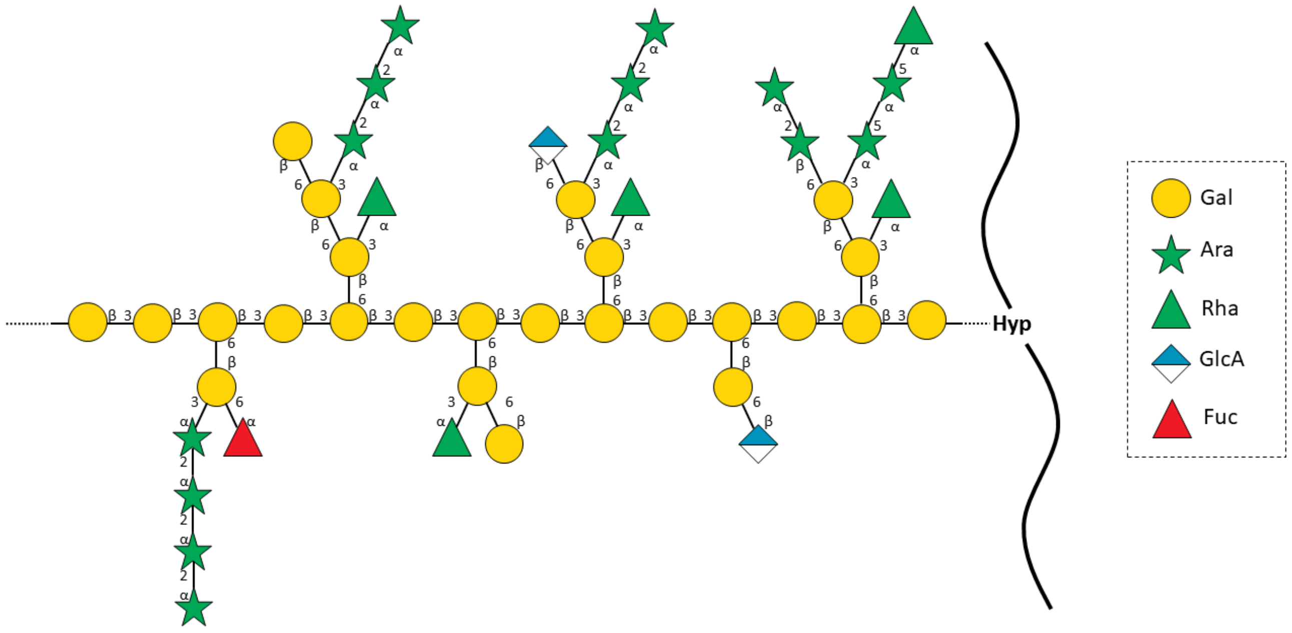


**Figure S1.** Proposed structure for the carbohydrate moiety of AGPs from *Azolla filiculoides*. The proposal was derived from the compositional, immunocytochemical, and linkage-type analyses of AGP and hydrolysed AGPs.

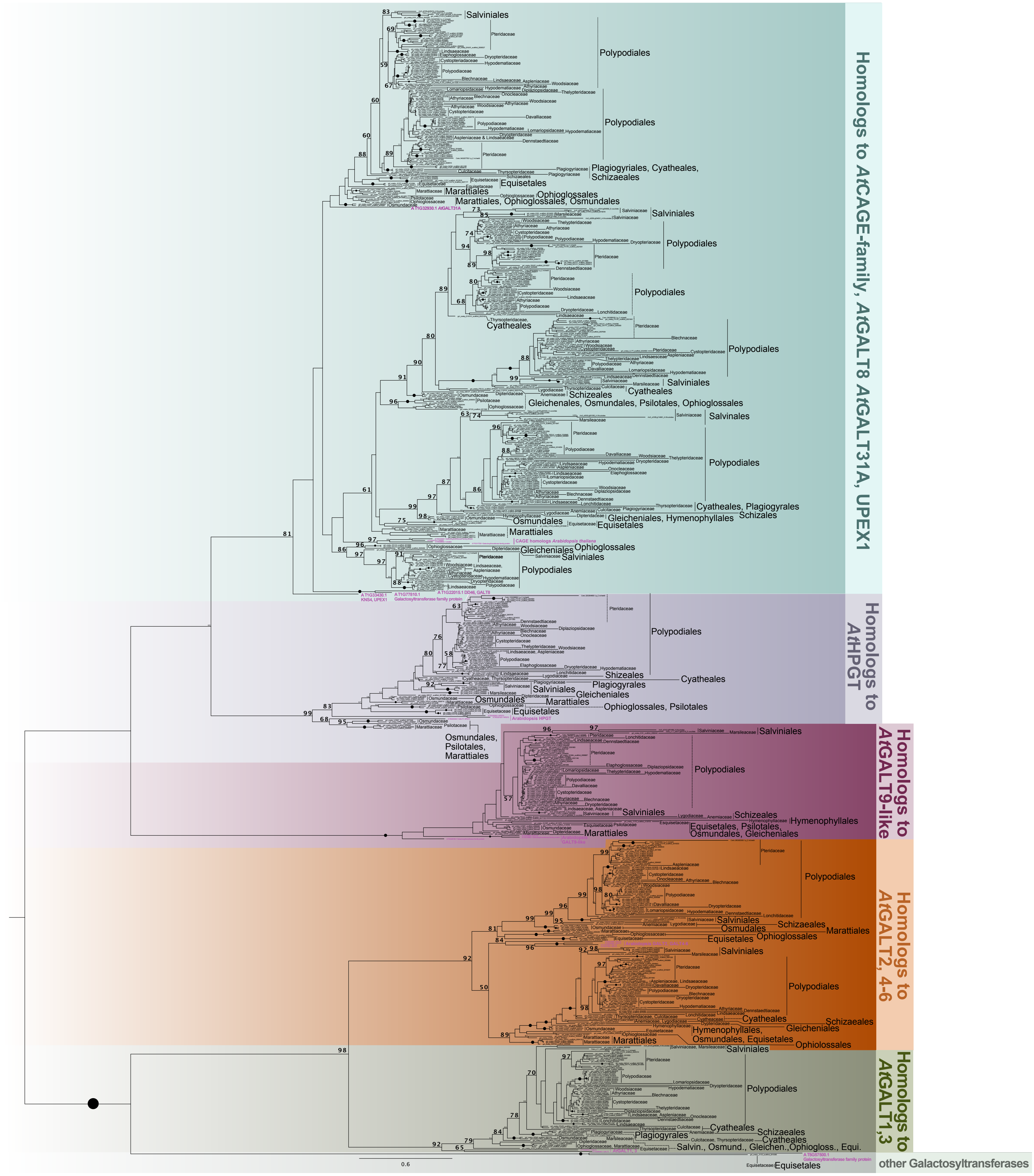


**Figure S2.** Phylogeny of GT31 enzymes throughout the different groups of ferns. The phylogeny was build using the model JTT+I+G4 and 100 bootstrap replicates. Bootstrap values ≤50 are not shown. Bootstrap values of 100 are indicated by black circles. Fern families and orders are indicated next to the phylogeny; here multiple accessions belonged to the same family or genus are group by lines. Solid lines indicate phylogenetically supported clades and dashed lines indicate that the accessions belonging to the same taxonomic association do either not group in a clade or that this clade has no phylogenetic support. *Arabidopsis thaliana* sequences are indicated in bold and purple. The phylogeny is midpoint rooted.

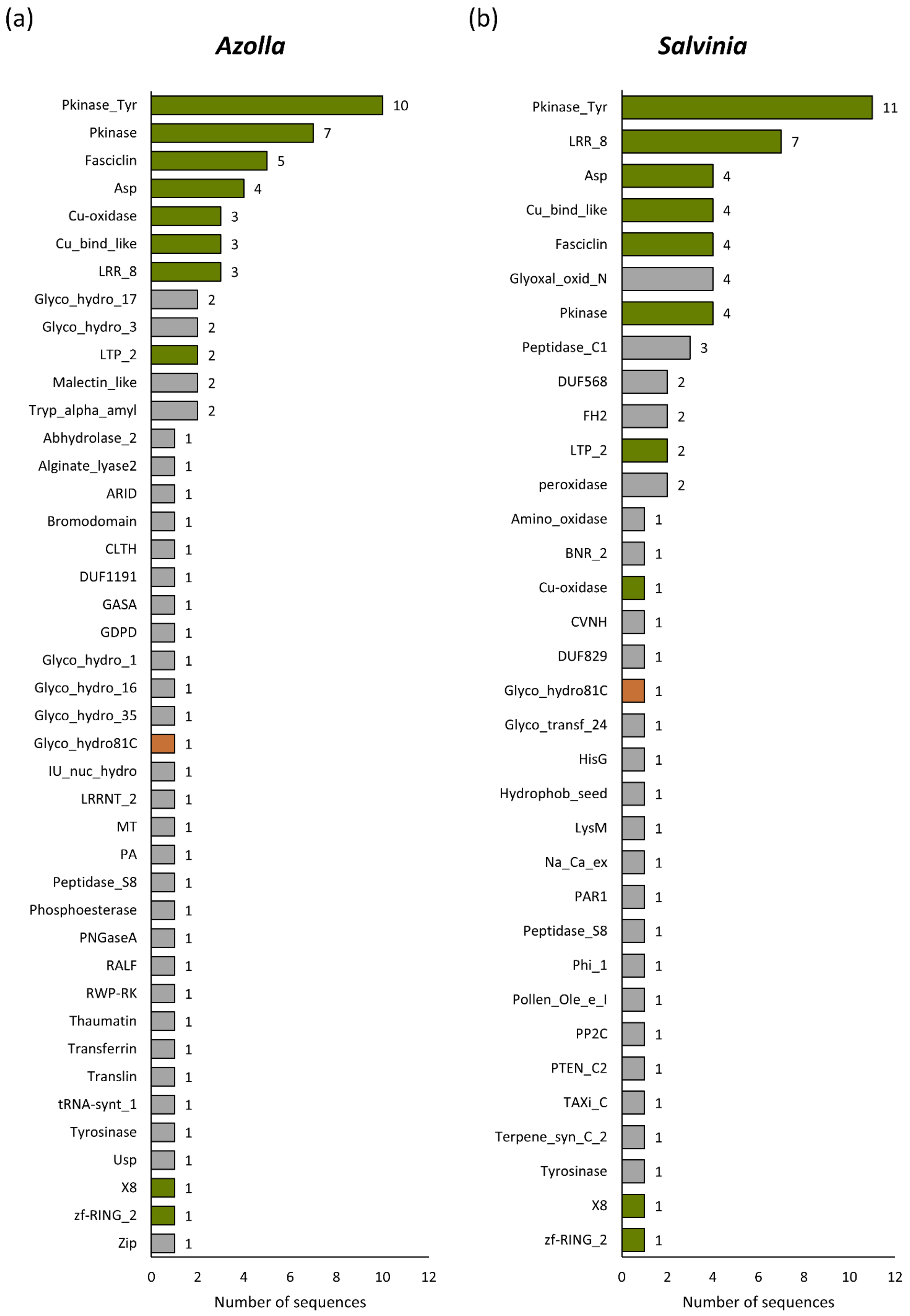
(a)
Azolla
Pkinase_Tyr 10
Pkinase 7
Fasciclin 5
Asp 4
Cu-oxidase 3
Cu_bind_like 3
LRR_8 3
Glyco_hydro_17 2
Glyco_hydro_3 2
LTP_2 2
Malectin_like 2
Tryp_alpha_amyl 2
Abhydrolase_2 1
Alginate_lyase2 1
ARID 1
Bromodomain 1
CLTH 1
DUF1191 1
GASA 1
GDPD 1
Glyco_hydro_1 1
Glyco_hydro_16 1
Glyco_hydro_35 1
Glyco_hydro81C 1
IU_nuc_hydro 1
LRRNT_2 1
MT 1
PA 1
Peptidase_S8 1
Phosphoesterase 1
PNGaseA 1
RALF 1
RWP-RK 1
Thaumatin 1
Transferrin 1
Translin 1
tRNA-synt_1 1
Tyrosinase 1
Usp 1
X8 1
zf-RING_2 1
Zip 1
0 2 4 6 8 10 12
Number of sequences
(b)
Salvinia
Pkinase_Tyr 11
LRR_8 7
Asp 4
Cu_bind_like 4
Fasciclin 4
Glyoxal_oxid_N 4
Pkinase 4
Peptidase_C1 3
DUF568 2
FH2 2
LTP_2 2
peroxidase 2
Amino_oxidase 1
BNR_2 1
Cu-oxidase 1
CVNH 1
DUF829 1
Glyco_hydro81C 1
Glyco_transf_24 1
HisG 1
Hydrophob_seed 1
LysM 1
Na_Ca_ex 1
PAR1 1
Peptidase_S8 1
Phi_1 1
Pollen_Ole_e_I 1
PP2C 1
PTEN_C2 1
TAXi_C 1
Terpene_syn_C_2 1
Tyrosinase 1
X8 1
zf-RING_2 1
0 2 4 6 8 10 12
Number of sequences

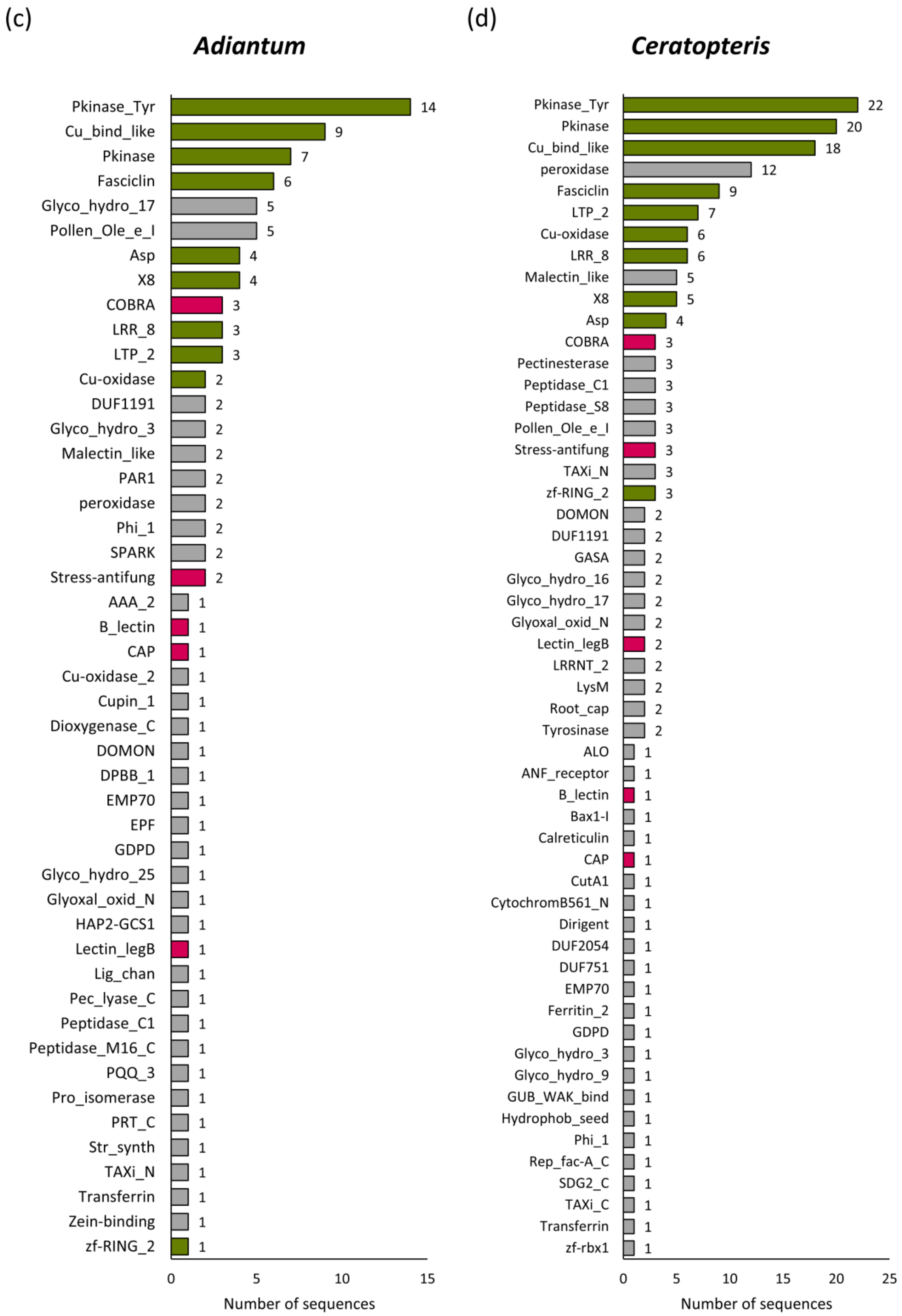

(c)
Adiantum
Pkinase_Tyr 14
Cu_bind_like 9
Pkinase 7
Fasciclin 6
Glyco_hydro_17 5
Pollen_Ole_e_I 5
Asp 4
X8 4
COBRA 3
LRR_8 3
LTP_2 3
Cu-oxidase 2
DUF1191 2
Glyco_hydro_3 2
Malectin_like 2
PAR1 2
peroxidase 2
Phi_1 2
SPARK 2
Stress-antifung 2
AAA_2 1
B_lectin 1
CAP 1
Cu-oxidase_2 1
Cupin_1 1
Dioxygenase_C 1
DOMON 1
DPBB_1 1
EMP70 1
EPF 1
GDPD 1
Glyco_hydro_25 1
Glyoxal_oxid_N 1
HAP2-GCS1 1
Lectin_legB 1
Lig_chan 1
Pec_lyase_C 1
Peptidase_C1 1
Peptidase_M16_C 1
PQQ_3 1
Pro_isomerase 1
PRT_C 1
Str_synth 1
TAXi_N 1
Transferrin 1
Zein-binding 1
zf-RING_2 1
0 5 10 15
Number of sequences
(d)
Ceratopteris
Pkinase_Tyr 22
Pkinase 20
Cu_bind_like 18
peroxidase 12
Fasciclin 9
LTP_2 7
Cu-oxidase 6
LRR_8 6
Malectin_like 5
X8 5
Asp 4
COBRA 3
Pectinesterase 3
Peptidase_C1 3
Peptidase_S8 3
Pollen_Ole_e_I 3
Stress-antifung 3
TAXi_N 3
zf-RING_2 3
DOMON 2
DUF1191 2
GASA 2
Glyco_hydro_16 2
Glyco_hydro_17 2
Glyoxal_oxid_N 2
Lectin_legB 2
LRRNT_2 2
LysM 2
Root_cap 2
Tyrosinase 2
ALO 1
ANF_receptor 1
B_lectin 1
Bax1-I 1
Calreticulin 1
CAP 1
CutA1 1
CytochromB561_N 1
Dirigent 1
DUF2054 1
DUF751 1
EMP70 1
Ferritin_2 1
GDPD 1
Glyco_hydro_3 1
Glyco_hydro_9 1
GUB_WAK_bind 1
Hydrophob_seed 1
Phi_1 1
Rep_fac-A_C 1
SDG2_C 1
TAXi_C 1
Transferrin 1
zf-rbx1 1
0 5 10 15 20 25
Number of sequences

(e)

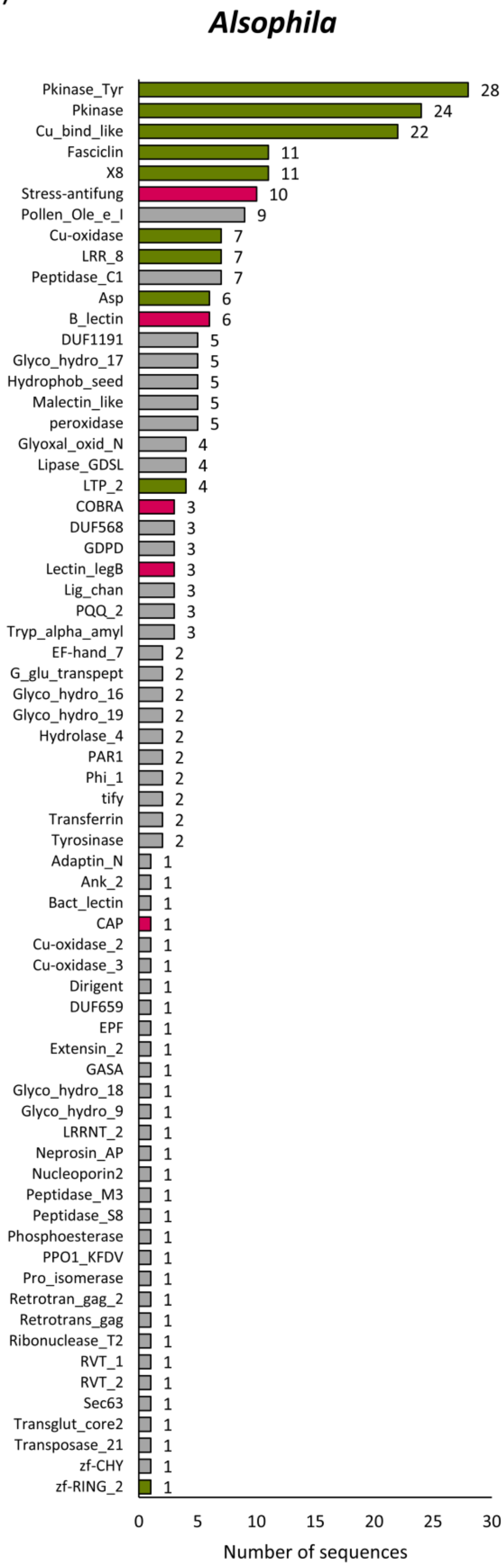


**Figure S3:** Chimeric AGPs of all investigated fern genomes ranked by decreasing numbers. If multiple domains occur in one protein sequence, the longest was counted. The color code of Figure 5 was used for common domains. (a) *Azolla* ,(b) *Salvinia*, (c) *Adiantum*, (d) *Ceratopteris*, (e) *Alsophila*.
